# stDyer enables spatial domain clustering with dynamic graph embedding

**DOI:** 10.1101/2024.05.08.593252

**Authors:** Ke Xu, Yu Xu, Zirui Wang, Xin Zhou, Lu Zhang

**Affiliations:** Department of Computer Science, Hong Kong Baptist University, Kowloon Tong, Hong Kong; Department of Biomedical Engineering, Vanderbilt University, 2301, Vanderbilt Place, Nashville, 37235, Tennessee, USA

**Keywords:** Spatially resolved transcriptomics, Spatial domain clustering, Dynamic graphs, Deep learning

## Abstract

Spatially resolved transcriptomics (SRT) data provide critical insights into gene expression patterns within tissue contexts, necessitating effective methods for identifying spatial domains. Traditional clustering techniques often overlook spatial information, leading to disjointed domains. Current computational approaches integrate spatial information but still face challenges in recognizing domain boundaries, scalability, and the need of independent clustering steps. We introduce stDyer, an end-to-end deep learning framework designed for spatial domain clustering in SRT data. stDyer combines a Gaussian Mixture Variational AutoEncoder (GMVAE) with graph attention networks (GATs) to simultaneously learn deep representations and perform clustering for units. A unique feature of stDyer is the dynamic graphs it adopts, which adaptively links units based on Gaussian Mixture assignments in the latent space, thereby improving spatial domain clustering and producing smoother domain boundaries. Additionally, stDyer’s mini-batch neighbor sampling strategy facilitates scalability to large datasets and enables multi-GPU training. Benchmarking against state-of-the-art tools across various SRT technologies, stDyer demonstrates superior performance in spatial domain clustering, multi-slice analysis, and large-scale dataset handling.

## Introduction

Identifying spatial domains is a vital task for analyzing spatially resolved transcriptomics (SRT). These domains represent distinct areas where the units involved (either spots or cells, depending on the SRT technologies) show similar gene expression patterns. Exploring these domains is crucial to understanding the spatial organization of gene activity [1, 2]. Traditional methods for clustering or community detection, such as K-means and the Leiden algorithm, typically rely only on gene expression data and disregard spatial information, often resulting in spatial domains that lack spatial continuity. To address this issue, various algorithms have been developed to incorporate both gene expression and spatial coordinates. These methods can be broadly classified into probabilistic approaches and deep learning models.

In probabilistic approaches, HMRF [3] groups units with similar gene expression profiles and takes into account their spatial coordinates using a Hidden Markov Random Field. HMRF employs the expectation-maximization algorithm and Gaussian density function to estimate model parameters. However, estimating a large number of parameters in the variable precision matrix across clusters can lead to unstable results.BayesSpace [4] employs a *t* -distributed error model to generate more robust spatial domains. It improves HMRF by utilizing Markov Chain Monte Carlo for parame-ter inference, allowing for more efficient exploration in the parameter space. BASS [5] implements a Bayesian hierarchical framework to simultaneously identify spatial domains and their featured cell types across multiple samples.

Deep learning models employ graph convolutional networks (GCNs) [6] on a spatial graph constructed using spatial coordinates of units, such as the K-nearest neighbor (KNN) graph. They group unit embeddings using unsupervised clustering methods to recognize spatial domains. SpaGCN [7] builds a graph by integrating the spatial coordinates and histological features of the units and applies GCNs to aggregate information from the neighboring units on this graph. STAGATE [8] introduces graph attention networks (GATs) to a spatial graph to capture intricate spatial dependencies on gene expression and incorporates a cell type-aware module from pre-clustered gene expressions. GraphST [9] enhances the graph embeddings of units by adopting a graph self-supervised contrastive learning model that considers both gene expression profiles and spatial information.

Although the existing methods have been successfully applied to identify spatial domains for various tissues and species, many studies have explored that their performance is still unsatisfactory [1, 10, 11]. All these methods rely on a fixed spatial graph and assume that a unit is derived from the same spatial domain as its neighboring units. However, this assumption may not hold true for units located at the boundaries of spatial domains, where neighboring units may belong to different domains. The mis-classification of those boundary units may also extend its influence to units within the domain. Additionally, most existing deep learning models for spatial domain clustering [8, 12–14] are not designed to be end-to-end, resulting in suboptimal performance as they necessitate an independent clustering step. In addition, the application of existing tools to multi-slice and large-scale datasets poses another challenge with respect to scalability [15]. It has been observed that probabilistic approaches often require extensive processing time for large-scale datasets due to the absence of GPU acceleration, while deep learning models face limitations in graphics memory (Figure 7e) because they need to load the entire spatial graph into memory. There are currently no methods tailored for spatial domain clustering that can effectively address all of these three issues simultaneously.

We introduce stDyer, an end-to-end and scalable deep spatial domain clustering approach on SRT data. stDyer employs a Gaussian Mixture Variational AutoEncoder [16–18] (GMVAE) with GAT [19] and graph embedding in the latent space. Instead of using an independent clustering step, stDyer enables deep representation learning and clustering from Gaussian Mixture Models (GMMs) simultaneously. stDyer also introduces dynamic graphs to include more edges to a KNN spatial graph. These new edges link a unit to others that belong to the same Gaussian Mixture as it does and increase the likelihood that units at the domain boundaries establish connections with others belonging to the same spatial domain. Moreover, stDyer introduces mini-batch neighbor sampling strategy to enable its application to large-scale datasets. To our knowledge, stDyer is also the first method that could enable multi-GPU training for spatial domain clustering. We compared stDyer with seven state-of-the-art tools on four different SRT technologies, including 10x Visium, osmFISH, STARmap and Stereo-seq. stDyer demonstrated excellent performance on these technologies and produced smooth domain boundaries. In the ablation studies, we demonstrated that dynamic graphs could substantially improve spatial domain clustering when compared to the utilization of a KNN spatial graph (stDyer(KNN)). stDyer could also be extended to jointly analyze SRT data from multiple slices and scale up to large-scale datasets.

## Results

### Workflow of stDyer

stDyer employs GMVAE with GAT and graph embedding on dynamic graphs to improve spatial domain clustering using SRT data (Figure 1). We constructs a KNN spatial graph using the spatial coordinates of units within SRT data. This graph, together with gene expression profiles, serves as input for the GMVAE (Supplementary Figure S5b). The encoder layer of the GMVAE is structured as GATs (Supplementary Figure S5a) to consider the spatial graph structure (**Methods**). Besides GATs, stDyer also considers the graph structure in the latent space by encouraging the gene expression of a unit to be reconstructed by its neighbors in the graph. GMVAE assumes unit embeddings follow GMMs in the latent space, which could be used to generate temporary spatial domain labels for all units at each epoch. In the first epoch using dynamic graph for training, stDyer modifies the KNN spatial graph by connecting unit *i* to extra neighboring units (12 for the 10x Visium technology and 4 for the other technologies), where these additional units and unit *i* must share the same temporary spatial label (**Methods**). This is the initial dynamic graph we generated. Starting from the next training epoch, dynamic graphs are continuously updated by substituting the extra units based on temporary spatial labels from GMMs. stDyer includes a post-processing module to remove outliers and smooth spatial domain boundaries by jointly considering the predicted spatial labels of all units (**Methods**). stDyer also applies integrated gradient (IG) analysis (**Methods**) to identify spatially variable genes (SVGs) with spatially correlated expression.

**Fig. 1:**
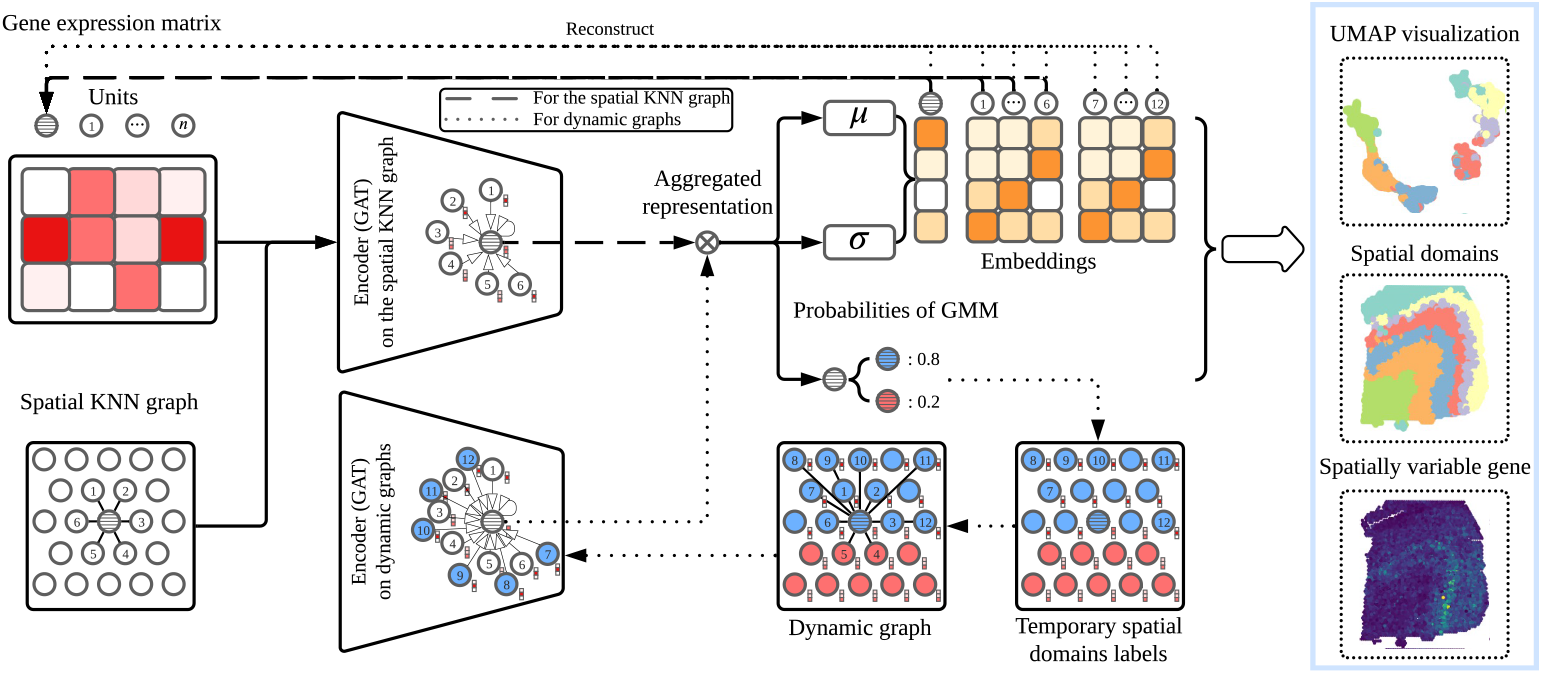
The workflow of stDyer. stDyer accepts gene expression profiles and a KNN spatial graph (K=6, units 1 to 6 for the shallow unit) from SRT data as inputs. GMVAE with GATs generates the unit embeddings and the probabilities of a unit belonging to Gaussian mixtures. The parameters for GMMs (*μ* and *σ*) and temporary spatial domain labels of units are estimated by maximizing the log-likelihood of the marginal distribution across all units (**Methods**). In the latent space, stDyer encourages the embedding of a unit to be reconstructed by its neighboring units in a KNN (dynamic) graph (e.g., 6 neighbors for a KNN spatial graph and 12 neighbors for a dynamic graph in this example). The temporary spatial domain label (red and blue) for each unit is generated by GMMs after each epoch. At the first training epoch, the KNN spatial graph is updated by connecting each target unit (e.g., shallow unit) to 6 additional units (e.g., units 7 to 12), where all these 7 units should share the same temporary spatial domain label. This dynamic graph and temporary spatial domain labels will be continuously updated after each epoch. stDyer also enables the identification of spatially variable genes using integrated gradient analysis.

### stDyer improves spatial domain identification on the DLPFC dataset from 10x Visium technology

We evaluated the performance of stDyer using the human dorsolateral prefrontal cortex (DLPFC) dataset [1] generated by 10x Visium technology. This dataset includes 12 slices, with unit numbers ranging from 3,460 to 4,789. Each slice has 5 or 7 annotated spatial domains (Supplementary Figure S1). We benchmarked stDyer with 8 state-of-the-art methods for spatial domain clustering: HMRF, SpaGCN, BayesSpace, SEDR, DeepST, BASS, STAGATE, and GraphST based on adjusted rand index (ARI; **Methods**). We evaluated these methods on all 12 slices (Figure 2f) and observed stDyer achieved an average ARI of 0.612, which was significantly higher than the second-best method GraphST (average ARI=0.538, p-value=0.014, Wilcoxon rank-sum test). We also performed an ablation study to examine the efficacy of using dynamic graphs in stDyer. Our results indicated that stDyer (average ARI=0.612, p-value=0.002, Figure 2f) substantially outperformed stDyer (KNN) (average ARI=0.486).

**Fig. 2:**
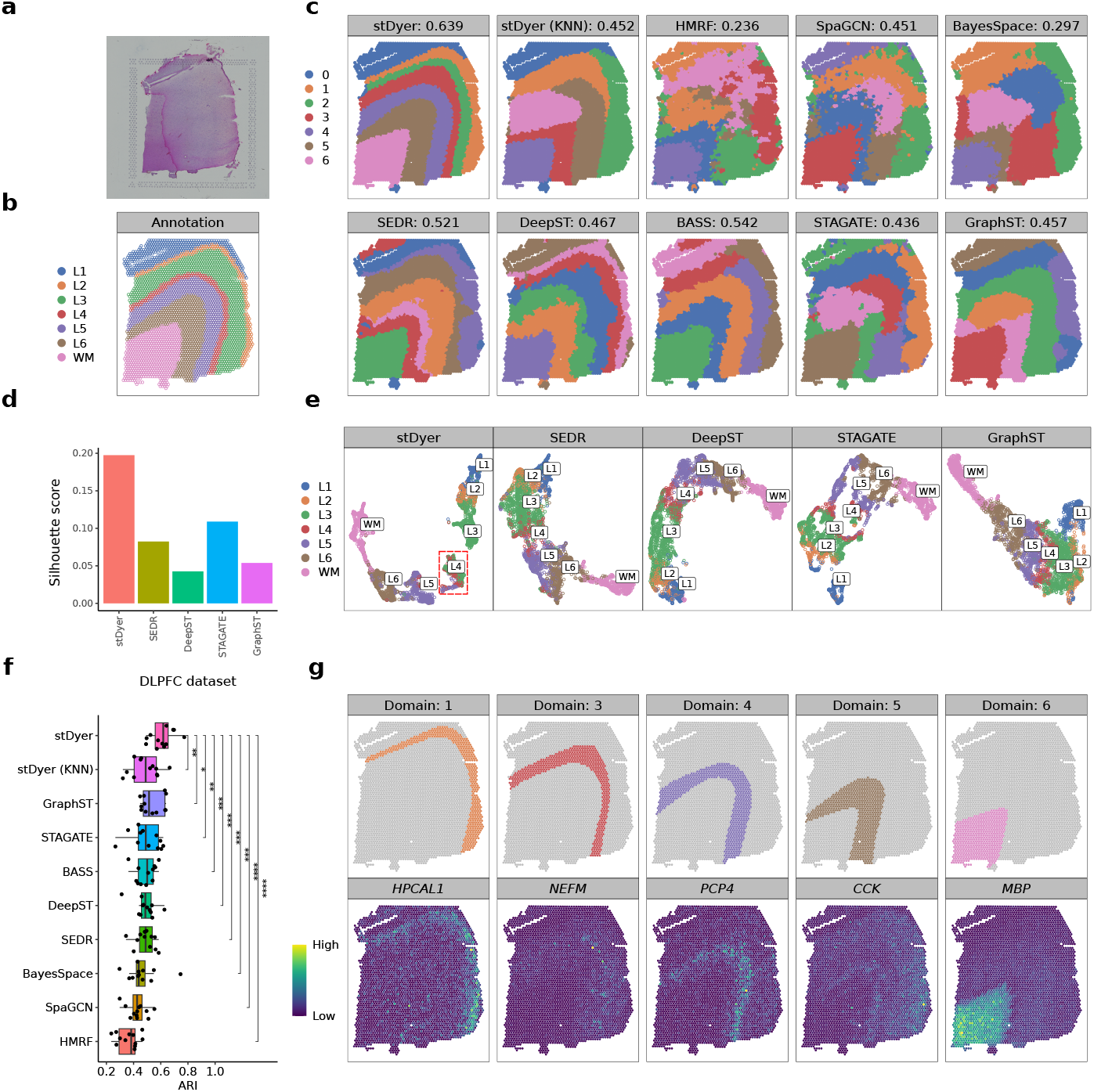
The performance of stDyer on 12 slices from the DLPFC dataset generated by 10x Visium technology. (**a**) The histology image of slice 151674 of the DLPFC dataset.(**b**) The visualization of spatial domain annotation of slice 151674. (**c**) Spatial domain clustering and ARI scores of different methods on slice 151674. (**d**) The Silhouette scores of the five methods that can generate unit embeddings in the latent space.(**e**) UMAP plots of unit embeddings generated by the five methods in **d**. (**f**) The boxplot of ARIs of different methods on all 12 slices. The p-values were computed by the Wilcoxon rank-sum test (**\*\*\*\***: p-value*<*0.0001; **\*\*\***: p-value*<*0.001; **\*\***: p-value*<*0.01; **\***: p-value*<*0.05). (**g**) Visualization of SVG expression values (bottom) and their associated spatial domains (top). ARI: adjusted rand index. stDyer (KNN): stDyer with a KNN spatial graph.

By visualizing slice 151674 as an example (Figure 2a), only stDyer (Figure 2c) could identify six laminar layers and white matter (WM) in the annotation (Figure 2b). Furthermore, the embeddings from stDyer exhibited continuity and adhered to the adjacency pattern in the annotation from L1 (domain 0) to WM (domain 6) (Figure 2e). There were five methods that could generate unit embeddings (stDyer, SEDR, DeepST, STAGATE and GraphST). Among these five methods, stDyer produced the highest Silhouette score (Figure 2d) and it was the only one that could separate out L4 domain (highlighted by the red rectangle in Figure 2e). The other methods struggled to distinguish between L2 to L6, e.g., GraphST failed to separate L4 and L5 domains (Figure 2c). Similar patterns were observed on the other slices (Supplementary Figure S1).

We selected 50 SVGs with the highest IG values for each domain (**Methods**, Supplementary Table S2 and Supplementary Figure S2) and found some of them were reported as marker genes of cortex layers. For example, a previous study identified *HPCAL1, PCP4*, and *MBP* as marker genes of L2, L5, and WM [1] (Figure 2g), respectively. The same study also revealed *CCK* could distinguish L2, L3, and L6 from the other layers. We found that *NEFM* could be a marker gene of L4.

### stDyer supports multi-slice spatial domain clustering

We divided 12 slices from the DLPFC dataset into three slice sections according to their spatial distance of the third dimension, thus each section includes four adjacent slices (Figure 3d, Table S5) without obvious batch effects (Figure 3e with slice ids) [1]. The KNN spatial graph was constructed using three-dimensional coordinates of units (**Methods**), which could include neighboring units from the same and adjacent slices. By considering four slices (151507, 151508, 151509 and 151510) from a section jointly, we observed multi-slice stDyer (stDyer (M), ARI=0.572) significantly surpassed single-slice stDyer on slice 151510 (stDyer (S), ARI=0.496, Figure 3c). We also compared stDyer(M) with three methods (GraphST(M), STAGATE(M) and SEDR(M)) that also supported multi-slice clustering and generated unit embeddings. The stDyer(M) achieved an average ARI of 0.650 for all three sections, substantially outperforming the second-best method, GraphST, which had an average ARI of 0.540 (p-value=0.019, Figure 3a). We also observed stDyer(M) provided more accurate unit embeddings of slender domains including L2, L4, L5 and L6 (Figure 3e). Both Silhouette scores (Figure 3b) and UMAP visualization (Figure 3e) suggested that unit embeddings from stDyer(M) were much more consistent with the spatial domain annotation.

**Fig. 3:**
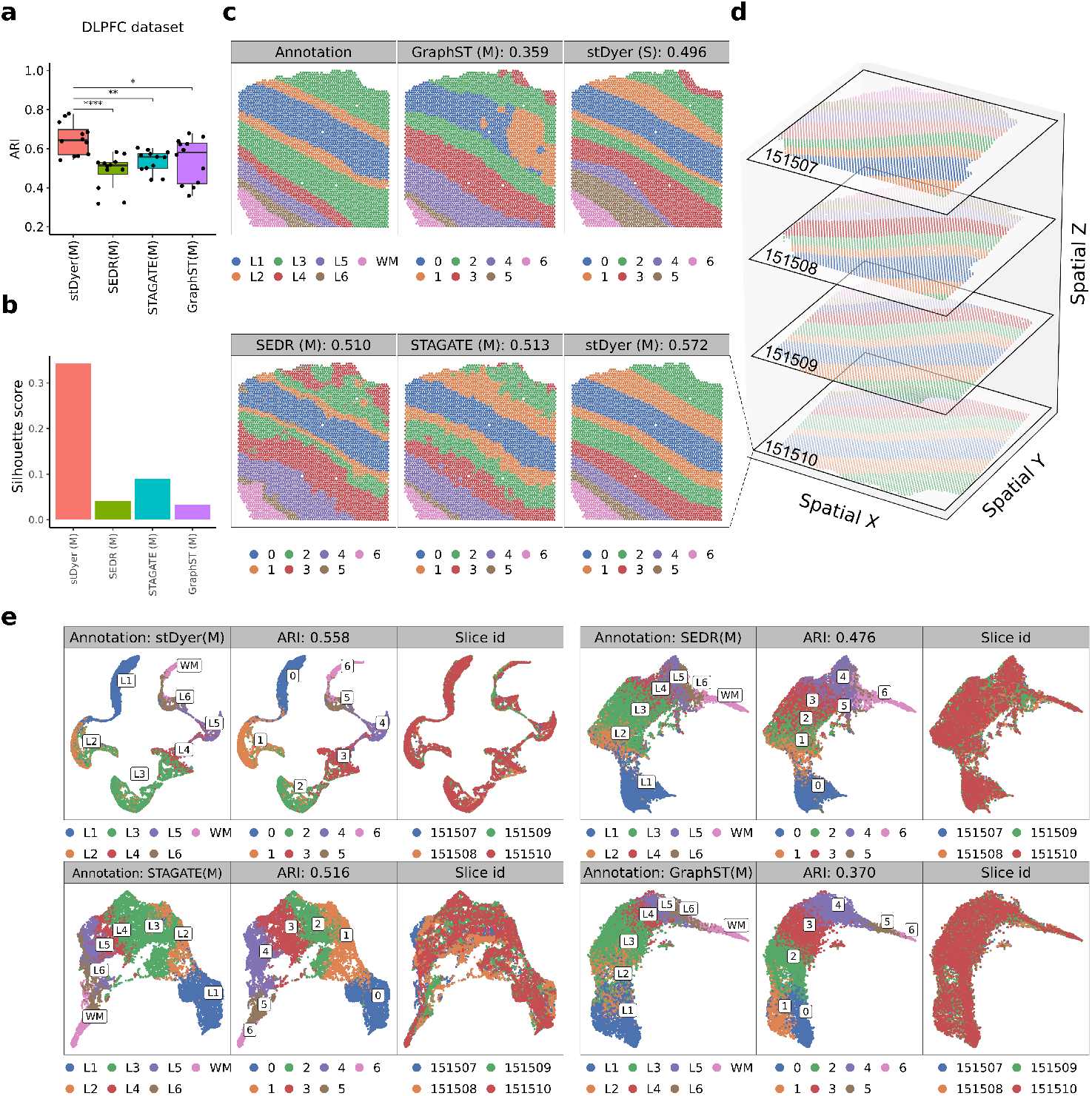
The performance of stDyer on the DLPFC dataset by jointly considering multiple slices. (**a**) Comparison of four methods that support spatial domain clustering on multiple slices. The p-values were computed by the Wilcoxon rank-sum test (**\*\*\*\***: p-value*<*0.0001; **\*\***: p-value*<*0.01; **\***: p-value*<*0.05). (**b**) The Silhouette scores of the four methods in (**a**). (**c**) Visualization of spatial domains and ARI scores for the four methods described in (**a**) and stDyer on the single slice (stDyer(S), slice 151510).stDyer(M): stDyer on four slices (101507, 151508, 151509, 151510).(**d**) The spatial organization of four slices in a section. (**e**) UMAP visualization by labeling units with the annotated spatial domains, predicted spatial domains and slice ids.

### stDyer recognizes both tumor and interface domains on a zebrafish tumor dataset from 10x Visium technology

We further applied stDyer to a zebrafish melanoma dataset from 10x Visium technology [11] to assess its ability to identify highly heterogeneous domains. This dataset comprises 2,179 units with 16,107 genes, including spatial domains from tumor, interface, and normal tissues. The normal tissue domains are further categorized into the brain, muscle, secondary muscle, skin and others. stDyer with dynamic graphs (ARI=0.616) was the only method that identified both tumor and interface domains (Figure 4a); it could also refine the boundary of the interface domain and recognize the secondary muscle domain compared to stDyer (KNN) (ARI=0.590). SpaGCN (ARI=0.445), BASS (ARI=0.418), SEDR (ARI=0.286) and DeepST (ARI=0.452) could correctly identify the interface domain, but they divided tumor domains into more than one segment. BayesSpace (ARI=0.367), STAGATE (ARI=0.460) and GraphST (ARI=0.360) were unable to distinguish the interface domain from the surrounding units.

**Fig. 4:**
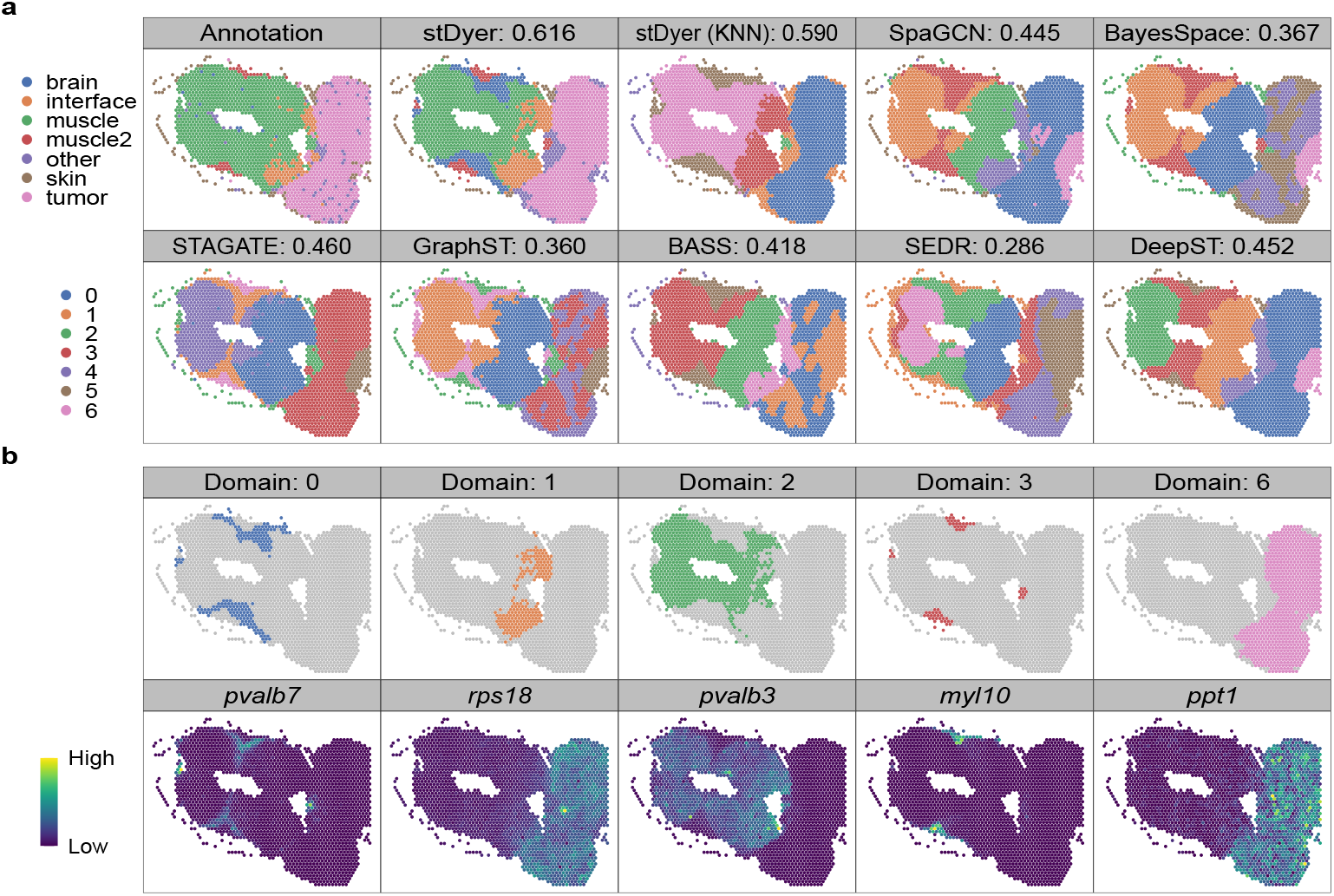
The performance of stDyer on a zebrafish tumor dataset from 10x Visium technology. (**a**) Visualization and ARIs of different methods for spatial domain clustering. (**b**) Visualization of SVG expression values (bottom) and their associated spatial domains (top). muscle2: secondary muscle.

Based IG information, we observed SVG expression values exhibited clear correlations with the corresponding spatial domains (Figure 4b, Supplementary Table S4 and Supplementary Figure S4). We found *ppt1* uniquely showed high expression profiles within the tumor domain (Figure 4b). Its homeotic gene *PPT1* has been found to be associated with tumor promotion and prognosis [20]. We noticed that the expression levels of *rps18* progressively decreased as the units transitioned from the tumor domain through the interface domain and into the muscle domain. This pattern indicated that *rps18* might play a role in tumor progression.

### stDyer generates spatial domains with smooth boundaries on the mouse cortex dataset from osmFISH

We analyzed a mouse cortex dataset comprising 4,839 units and 33 genes from osm-FISH [10] to evaluate the performance of stDyer on *in situ* hybridization technology. These units can be classified into 11 spatial domains, including hippocampus (HPC), internal capsule and caudoputamen (ICC), neocortical layer 1 (L1), neocortical layer 2-3 lateral (L2-3l), neocortical layer 2-3 medial (L2-3m), neocortical layer 3-4 (L3-4), neocortical layer 4 (L4), neocortical layer (L5), neocortical layer (L6), ventricle (VT), and white matter (WM). stDyer achieved the best performance with ARI of 0.733 (Figure 5a), which was higher than stDyer (KNN) (ARI=0.632) and all seven other state-of-the-art tools (the second-best tool BASS with ARI=0.573). stDyer (ARI=0.733), DeepST (ARI=0.561), BASS (ARI=0.573) and GraphST (ARI=0.373) could all generate notably smooth spatial domain boundaries (Figure 5a), whereas the remaining three methods preferred to label neighboring units with different domains.

**Fig. 5:**
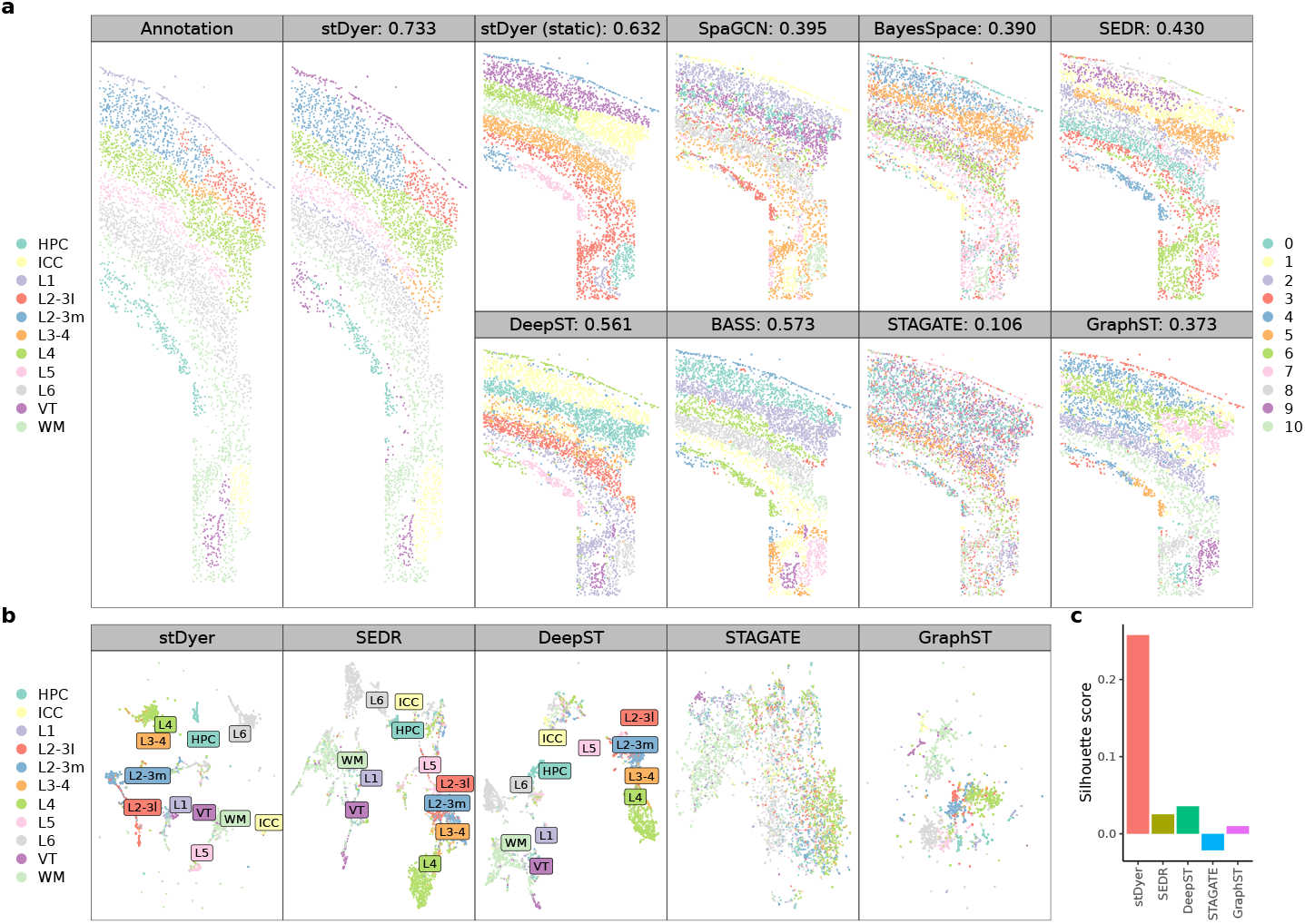
The performance of stDyer on a mouse cortex dataset from osmFISH. (**a**) Visualization and ARIs of different methods for spatial domain clustering. stDyer (KNN): stDyer with a KNN spatial graph. (**b**) UMAP visualization by labeling units with the annotated spatial domains. (**c**) The Silhouette scores for the five methods that provide unit embeddings.

Both DeepST and BASS incorrectly merged L2-3l and L2-3m domains in their results. stDyer generated highly separable unit embeddings from different domains (Figure 5b) and obtained the highest Silhouette score (Figure 5c). It was noted that stDyer generated unit embeddings of L2-3l domain as a line that could be easily distinguished from L2-3m (Figure 5b).

### stDyer detects clear laminar structures on the mouse cortex dataset from STARmap

stDyer was used to analyze a mouse cortex dataset [21] comprising 1,207 units and 1,020 genes from STARmap to evaluate the performance of stDyer on *in situ* sequencing technology. The units in this dataset were annotated to be from seven spatial domains, including corpus callosum (CC), hippocampus (HPC), neocortical layer 1 (L1), neocortical layer 2/3 (L2/L3), neocortical layer 4 (L4), neocortical layer 5 (L5), and neocortical layer 6 (L6). As shown in Figure 6a, stDyer showed superior performance (ARI=0.664) in predicting the laminar structure of the mouse cortex. The other tools demonstrated issues in identifying different layers. There were only a few cells annotated in the second and sixth domains predicted by BASS (ARI=0.645) and SEDR (ARI=0.584), which could be due to incorrect labeling of units from the domain boundaries. DeepST (ARI=0.531) and STAGATE (ARI=0.527) divided L2/L3 into two vertical sections. SpaGCN (ARI=0.443) failed to predict the hippocampus domain, while GraphST (ARI=0.465) was unable to predict L1 and divided L2/L3 erroneously. BayesSpace (ARI=0.219) exhibited poor performance on this dataset, yielding only four spatial domains with clear boundaries. We observed the unit embeddings from stDyer showed excellent separation between different spatial domains (Figure 6b) and achieved the highest Silhouette score (Figure 6c) compared to the other four methods. We observed that the IG analysis (Supplementary Table S3 and Supplementary Figure S3) also revealed SVGs associated with L2/3 and L6 (Figure 6d).

**Fig. 6:**
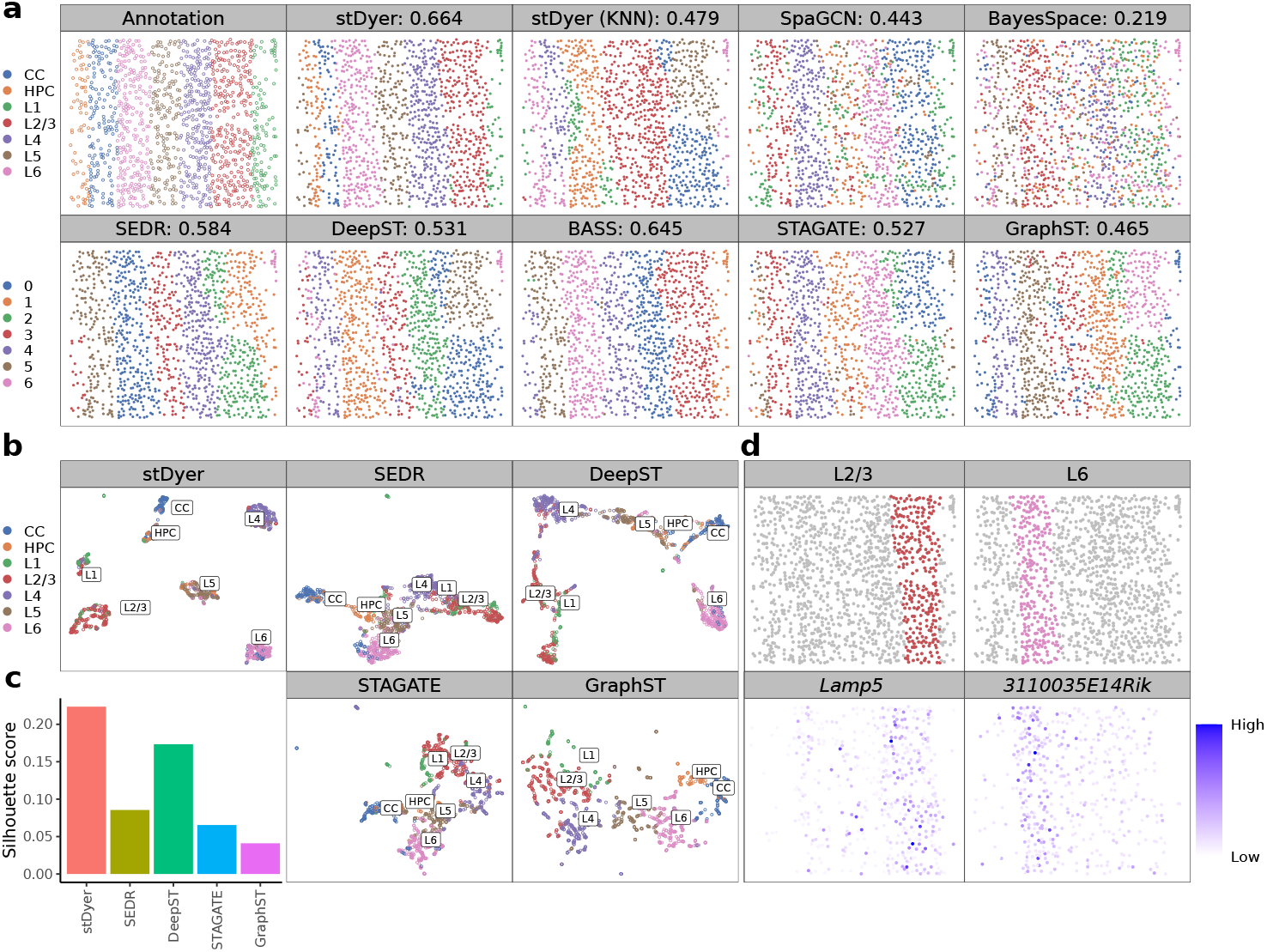
The performance of stDyer on a mouse cortex dataset from STARmap. (**a**) Visualization and ARIs of different methods for spatial domain clustering. stDyer detected clearer laminar structures and boundaries compared to the other tools. (**b**) UMAP visualization by labeling units with the annotated spatial domains. (**c**) The Silhouette scores for the five methods that generate unit embeddings. (**d**) Visualization of spatial domains (top) and their associated SVG expression values (bottom).

### stDyer can be scaled up to accommodate large-scale mouse embryo datasets from Stereo-Seq

Stereo-seq has been used to generate SRT data with 53 slices from 8 stages of mouse embryos [2]. We benchmarked stDyer with 7 existing methods (including SpaGCN, BayesSpace, SEDR, DeepST, BASS, STAGATE and GraphST) on 13 slices of E16.5 mouse embryo with unit numbers varying from 59,119 to 155,047 (Supplementary Table S1). We found DeepST, BASS and STAGATE failed to analyze the slice with the smallest number of units (Slice ID: 1) due to insufficient graphics memory (*>*80GB for DeepST and STAGATE) or encountering segfault from C stack overflow (BASS), suggesting they could be inappropriate to be applied to Stereo-seq data. Among the five remaining methods, stDyer, SpaGCN, and BayesSpace were the only ones able to analyze all 13 slices. Furthermore, stDyer was the only method that could utilize GPU acceleration.

To further evaluate their performance on scalability, we sampled 100,000 to 500,000 units from the combined units from 13 slices. BayesSpace, however, encountered an error due to matrix size and could not process datasets larger than 200,000 units. SpaGCN’s memory consumption was significant, requiring 195GB, 762GB, and 2,048GB of host CPU memory for the datasets including 100,000, 200,000, and 300,000 units, respectively (Figure 7f). For the datasets with over 300,000 units, SpaGCN exhausted all 2,919 GB of available host CPU memory and was terminated by the operating system’s out-of-memory management. In contrast, stDyer was more memory-efficient, requiring only about 25GB of graphics memory for all five simulated datasets due to the implementation of mini-batch strategy. For each batch, stDyer only selects a small number of units as well as its neighbors to construct the spatial graph, avoiding loading the whole spatial graph from all units into the graphics memory (Figure 7g).

**Fig. 7:**
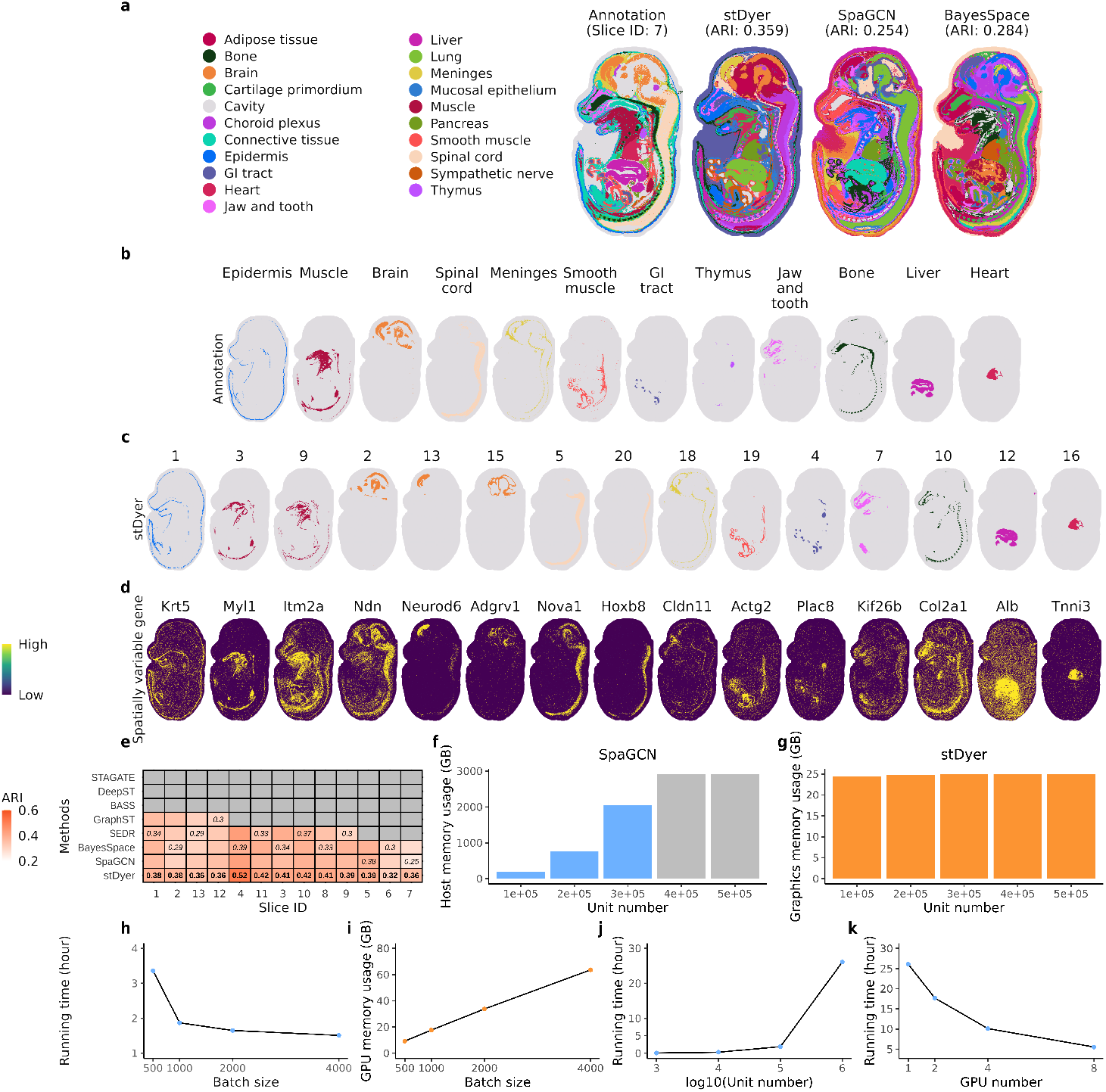
The performance of stDyer on 13 slices of E16.5 mouse embryos from Stereo-Seq. (**a**) Visualization of spatial domains predicted by stDyer, SpaGCN and BayesSpace on slice 7 (the largest slice). (**b**) Annotated spatial domains on slice 7. (**c**) Predicted spatial domains by stDyer on slice 7. The same color would be used for each mapping between an annotated domain and matched predicted domain(s). (**d**) Visualization of SVGs for the corresponding spatial domains in c. For each SVG, the gene expression values that are higher than the median values across all units are adjusted to the maximum values for better visualization. (**e**) Heatmaps of ARIs of 8 spatial domain clustering tools across 13 slices. The slice IDs are sorted in ascending order based on the unit numbers. The grey cells indicate methods fail to run on the corresponding slices. For each slice, we highlighted the maximum ARIs in bold and the minimum ARIs in italic. (**f**) Host CPU memory usage of SpaGCN. (**g**) Graphics memory usage of stDyer. (**h**) The impact of batch size on running time of stDyer to analyze 100,000 units. (**i**) The impact of batch size on GPU memory usage of stDyer to analyze 100,000 units. (**j**) The impact of the unit number on running time of stDyer. (**k**) The impact of GPU number on running time of stDyer to analyze 1,000,000 units.

We also investigated the impact of batch sizes and the number of units on the running time and GPU memory usage based on an NVIDIA A100 card (Figure 7h, i and j). We found that setting the batch size to 1,000 could significantly reduce the running time for processing 100,000 units (Figure 7h). It only required to occupy 17.64 GB of GPU memory (Figure 7i), which could even be satisfied by consumer GPUs like RTX 3090. By varying the total unit numbers (batch size=1,000, Figure 7j), we observed stDyer only required 1.87 and 26.11 hours to process 100,000 and 1,000,000 units, respectively. stDyer’s performance can be sped up further through the utilization of multiple GPU cards. Our findings indicated that by employing eight NVIDIA A100 cards, the analysis time for 1,000,000 units could be cut down to just 5.53 hours (Figure 7k).

stDyer also outperformed the other tools on spatial domain clustering across all 13 slices (Figure 7e). It achieved an average ARI of 0.393 (Supplementary Table S1) compared to 0.352 (SpaGCN) and 0.346 (BayesSpace), respectively. We also visualized the largest slice (Slice ID: 7) with 21 spatial domains, which could only be processed by stDyer, SpaGCN and BayesSpace. In this comparison, stDyer obtained the top ARI score of 0.359 (Figure 7a), which surpassed those achieved by SpaGCN (ARI=0.254) and BayesSpace (ARI=0.284). We also investigated if stDyer could identify major organs and tissues in mouse embryo accurately (Figure 7b and c). We found stDyer identified highly concordant spatial domains with annotations, and the SVGs (Figure 7d) of these domains also matched with the corresponding marker genes [22]or their homologous genes [23]. Specifically, the gene *Krt5* was identified for the Epidermis, *Myl1* and *Itm2a* for Muscle, *Nova1* and *Hoxb8* for Spinal cord, *Cldn11* for Meninges, *Actg2* for Smooth muscle, *Plac8* for GI tract and Thymus, *Kif26b* for Jaw and tooth, *Col2a1* for Bone, *Alb* for Liver and *Tnni3* for Heart. We also found stDyer separated the mouse brain into three distinct domains (domains 2, 13 and 15) and identified some SVGs associated with these domains that are functionally linked to the development of brain areas. For example, domain 13 was characterized by the presence of *Neurod6*, a gene known to be selectively active in certain deep-layer pyramidal neurons of the mouse brain [24]. In domain 15, the gene *Adgrv1* was found to be highly expressed, which is essential for the development of auditory functions[25]. Domain 2 worked in conjunction with the other two domains to define a particular brain area.

## Discussion

The SRT technologies have revolutionized our understanding of the spatial dynamics of cells and gene expression, providing an efficient way in investigating the intricate relationship between gene activity and its spatial distribution within tissue development or tumor microenvironment. Based on SRT data, spatial domains could be identified to investigate spatial heterogeneity of gene expression patterns and their functional implications. The performance of existing tools for spatial domain clustering is still unsatisfactory and it is challenging to accurately delineate the boundaries of spatial domains. In this study, we introduced stDyer as an end-to-end and scalable graph-based deep learning model for spatial domain clustering using dynamic graphs. Instead of using a fixed spatial graph, the edges in dynamic graphs would be updated based on units’ temporary spatial domain labels from GMVAE. Thus, there is a high chance that units sharing the same temporary spatial domain labels could be connected in dynamic graphs. We evaluated the performance of stDyer on different SRT technologies and tissues and proved it could substantially improve spatial domain clustering. The ablation studies also highlighted the superiority of the dynamic graphs over a KNN spatial graph.

Some SRT technologies, such as 10x Visium and Stereo-seq, are capable of producing histological images of the same tissue slice, which is expected to be beneficial for spatial domain clustering. We tried to concatenate image patch embeddings with gene expression profiles of the 151674 slice from the DLPFC dataset as input for our model. We cut the whole image into small patches for each unit and extracted the image feature embeddings using the pre-trained ResNet50 [26]. We found the ARI decreased to 0.237 (ARI without image: 0.639) and the predicted spatial domains were influenced much by noises in the histological image (Supplementary Figure S9). This suggested that image features could not guarantee to improve spatial domain clustering. To enhance clustering outcomes, the use of high-quality, biologically informative images or advanced image processing models will be desired.

Nowadays, advanced SRT technologies, such as Stereo-Seq, enable the measurement of a huge number of units for individual slices. Many graph-based approaches, such as STAGATE and GraphST, struggle to handle large spatial graphs that contain upwards of 100,000 units due to their reliance on a basic implementation of graph convolutional networks. These methods attempt to load the entire graph into the GPU graphics memory, which is inherently limited and unable to expand at the same rate as the growing number of units being measured. To address this issue, stDyer utilizes a mini-batch technique to efficiently analyze graphs with millions of units. Additionally,stDyer is also able to support multiple slices spatial domain clustering, where the slices could be aligned vertically (e.g., DLPFC) or horizontally adjacent (e.g., concatenated as a single large slice).

stDyer is an open-source tool with broad applicability for all existing SRT technologies. Although our experiments are limited to datasets produced by 10x Visium, osmFISH, STARmap, and Stereo-seq, stDyer is designed to handle datasets from any SRT technologies with gene expression profiles and spatial information. We have improved stDyer to enable the processing of multiple slices and incorporated both a mini-batch approach and multi-GPU support to efficiently handle the analysis of large-scale datasets. In the future, we would like to extend stDyer by incorporating histological images.

## Methods

### Data preprocessing

A SRT dataset comprises two primary components: one matrix representing the gene expression profiles of *m* genes in *n* units (***X*** ∈ ℝ_*n×m*_) and the other matrix containing two-dimensional spatial coordinates for *n* units (***C*** ∈ ℝ_*n×*2_). We performed data preprocessing on the gene expression matrix using functions provided by Scanpy [27]: 1. removed ERCC spike-ins; 2. removed units [scanpy.pp.filter cells(min counts=1)] and genes [scanpy.pp.filter genes(min counts=1)] with zero RNA counts; 3. selected top 3,000 highly variable genes [scanpy.pp.highly variable genes]; 4. library normalization [scanpy.pp.normalize total]; 5. logarithmic transformation [scanpy.pp.log1p]; 6. gene-wise z-score standardization [scanpy.pp.scale].

### Construct dynamic graphs

We established the initial spatial graph by employing a KNN approach (K = 4 for osmFISH, STARmap and Stereo-seq; K = 6 for 10x Visium due to its distinctive honeycomb structure) on the matrix ***C*** to link neighboring units based on Euclidean distance. During each training epoch of GMVAE, a temporary spatial domain label was assigned to each unit based on GMM (Supplementary Figure S6). Subsequently, additional neighbors (12 for 10x Visium and 4 for osmFISH, STARmap and Stereo-seq) for each unit were incorporated into the KNN spatial graph to connect units sharing identical spatial domain labels. For a given unit, the extra neighbors were chosen at random from other units that share the same temporary spatial domain label with it unless this unit’s closest 18 (for 10x Visium technology) or 8 (for other SRT technologies) spatial neighbors sharing the same label with it. In such a case, these closest units would be selected as the unit’s extra neighbors. In a dynamic graph, each unit has a total of 18 (10x Visium) or 8 neighbors (other SRT technologies).

### Preprocessing of SRT data from multiple slices

We divided 12 slices from the DLFPC dataset into three sections, each comprising four spatially contiguous slices, using the alignment method PASTE2 [28]. This method refines the existing two-dimensional spatial coordinates of units in the same slice to align them within a unified coordinate system. Given that each slice contains only two-dimensional spatial information, we introduced an additional artificial spatial coordinate of the third dimension to merge the four slices in the same section, thus enabling the construction of a three-dimensional spatial graph that spans across four slices. Assuming perfect overlap of the slices, this strategy allows each unit to identify 6 spatial neighbors from its own slice, as well as from the slices immediately above and below it, providing a total of 18 nearest neighbors for the three-dimensional spatial graph construction (Supplementary Note S8).

### The network structure of stDyer

To construct dynamic graphs, stDyer designs an end-to-end spatial domain clustering framework based on GMVAE with graph embedding (Supplementary Figure S5b). The framework includes an inference module (encoder) and a generative module (decoder). For the encoder, stDyer utilizes GATs (Supplementary Figure S5a) to aggregate gene expression profiles from the neighbors of a given unit in a spatial (dynamic) graph. Given a predefined number of clusters (*k*^cluster^), the expression profiles of the target unit *i* (***X***_*i*_) and its neighbors 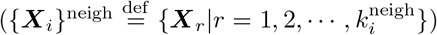, we formulate the inference component as follows:

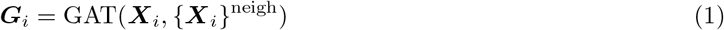

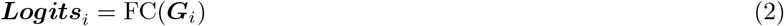

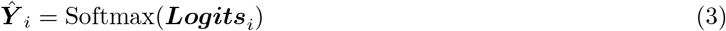

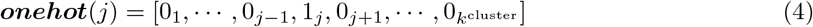

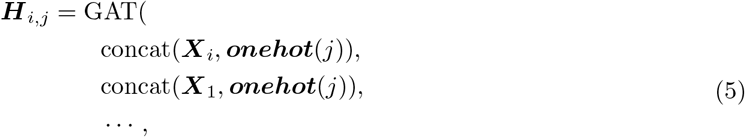

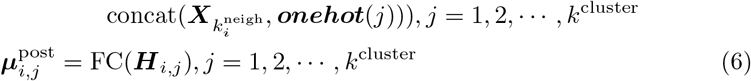

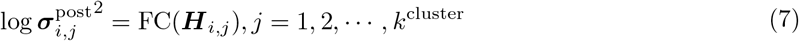

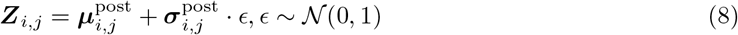

The aggregated gene expression from the neighboring units of *i* is represented by ***G***_*i*_, which is passed to a fully connected (FC) layer to generate ***Logits***_*i*_ to predict the temporary spatial domain label ***Ŷ*** _*i*_. The spatial domain labels of units are from the modes of GMM to which they are assigned. ***H***_*i*,*j*_ aggregates gene expression from neighboring units of unit *i* by assuming it belongs to a spatial domain *j*. ***H***_*i*,*j*_ is also used to derive parameters of the posterior distribution of the *j*th mode of GMM (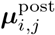 and 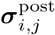) for 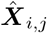. 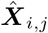 refers to the predicted ***X***_*i*_ from the *j*th mode of GMM. ***Z***_*i*,*j*_2*i*,*ji*,*ji*,*j* is the embedding of 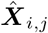 sampled from 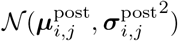 .

The decoder generates the parameters of the prior distribution of each compo-nent in GMM using ***onehot***(*j*). It is also required to reconstruct ***X***_*i*_ from ***Z***_*i*,*j*_. We formulate the generative module as follows:

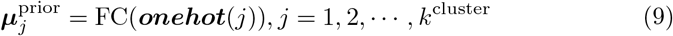

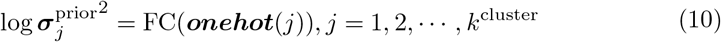

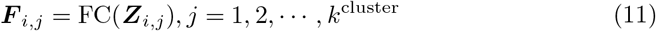

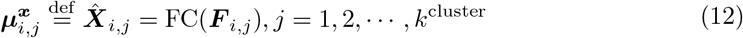

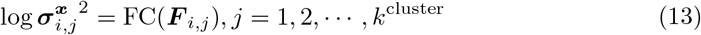

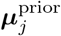 and 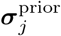 denote the mean and standard deviation of the prior distribution of the *j*th Gaussian mixture. 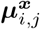 and 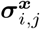 represent the mean and standard deviationof the distribution of 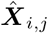 .

### Probabilistic models of modified GMVAE in stDyer

We modified the GMVAE in stDyer by replacing its encoder from FC layers to GAT and utilized *θ* and *ϕ* to represent the parameters involved during generative (decoder) and inference (encoder) processes. The generative process could be represented as follows:

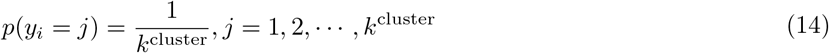

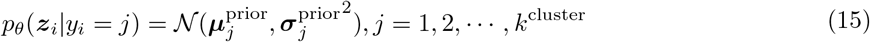

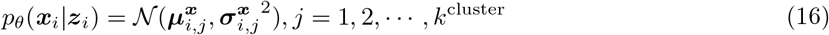

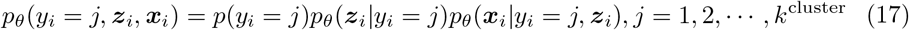

***x***_*i*_, *y*_*i*_ and ***z***_*i*_ are three random variables that denote the gene expression profile, spatial domain label and latent embedding from the *j*th mode of GMM of the target unit *i*. The inference process is formulated as follows:

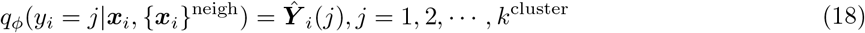

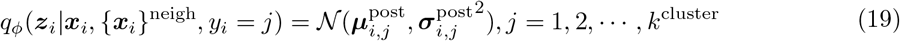

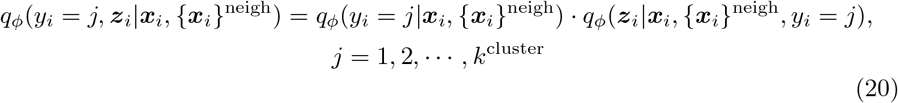

where ***Ŷ*** _*i*_(*j*) denotes the predicted probability that the target unit *i* belongs to the *j*th mode of GMM, {***x***_*i*_*}* ^neigh^ represents the gene expression profiles of the neighbors of unit *i*. The spatial domain assignment vector ***Ŷ*** _*i*_(*j*) is inferred by GAT in Equation (3).

### The objective function of stDyer

The objective function of stDyer is to maximize the log-likelihood of the marginal distribution across all involved *n* units:

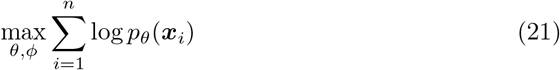

stDyer incorporates a smoothness constraint to encourage neighboring units to share the same spatial domain labels and have contiguous embedding.

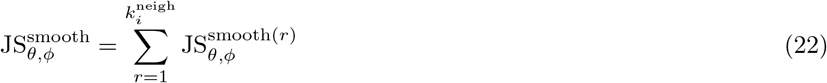

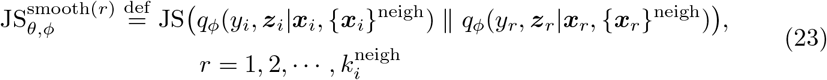

The objective function could be rewritten as:

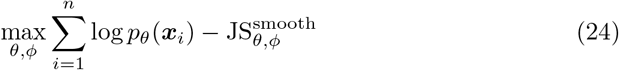

where JS denotes Jensen-Shannon divergence and ***x***_*r*_ refers to the gene expression profile of the neighboring unit *r* of unit *i*. The log-likelihood log *p*_*θ*_(***x***_*i*_) for the target unit *i* can be decomposed as:

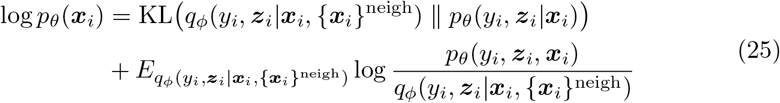

stDyer aims to reconstruct ***x***_*i*_ using its neighboring units ***x***_*r*_ to promote similarity in their spatial domain labels and latent embeddings. Thus, Equation (25) could be rewritten as:

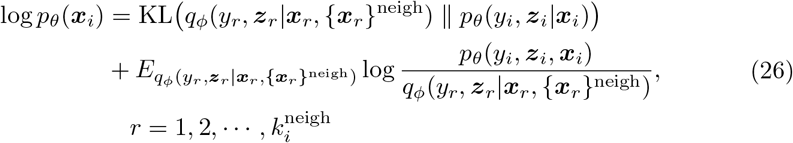

By taking the average of both sides of the two equations, Equation (25) and the mean of Equation (26) over *r* = 1,2, …, 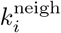, we derived:

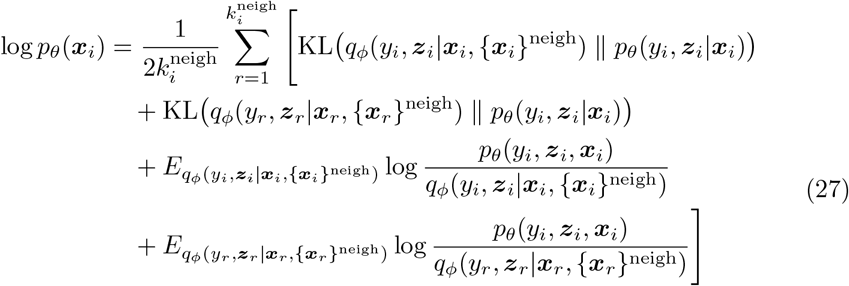

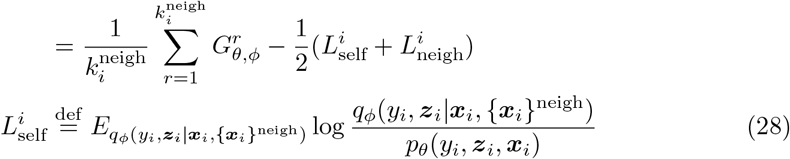

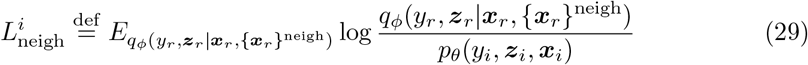

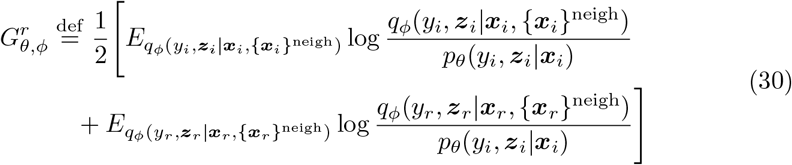

As 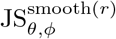 was the lower bound of 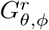 (Supplementary Note S6), the objective function Equation (24) could be reformatted with Equation (27) as:

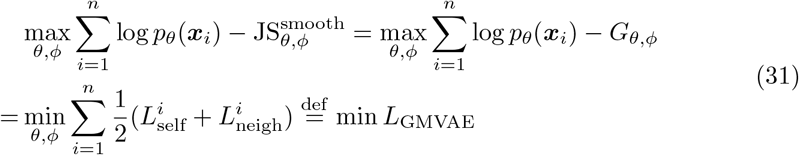

As defined in Equation (28) and Equation (29), 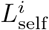 and 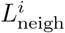 can be decomposed

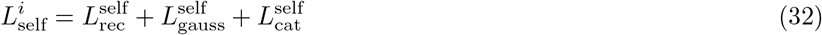

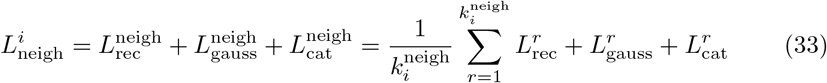

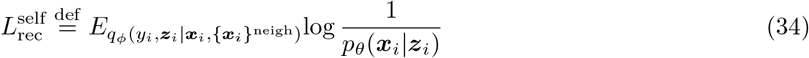

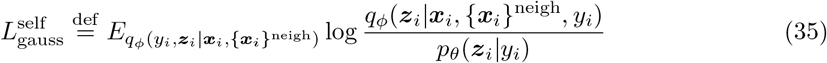

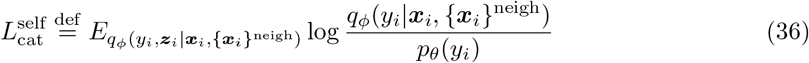

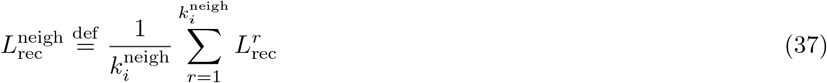

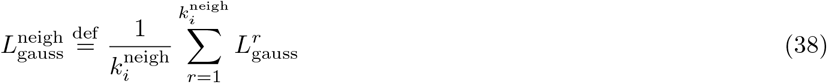

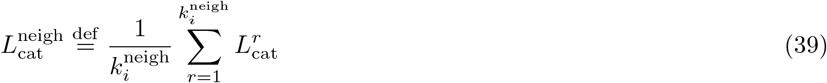

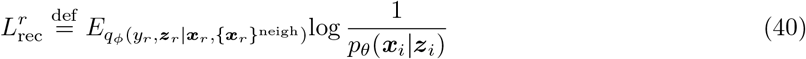

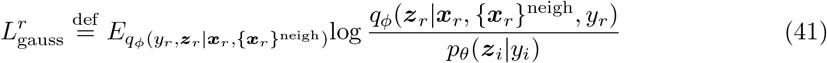

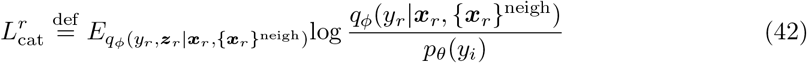

Here, 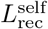 represents the loss when reconstructing the target unit from its own embedding, while 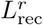 represents the loss when reconstructing the target unit from the embedding of its *r*th neighboring unit. Both 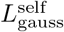 and 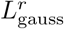 regularize theparameters of the posterior distribution to ensure consistency with those of the priordistribution in each mixture for the target unit *i* and its *r*th neighboring unit, respectively. The dim(***Z***_*i*,*j*_) denotes the number of dimensions of ***Z***_*i*,*j*_. Finally, 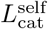 and 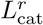 encourage the alignment between the prior distribution and the posterior distributionfor the predicted GMM assignment of the target unit and its *r*th neighbor unit.

Using previously defined generative and inference processes, 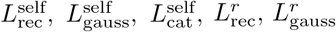 and 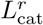 can be further expanded as:

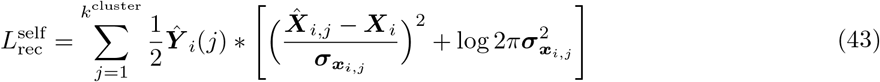

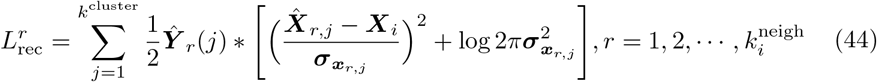

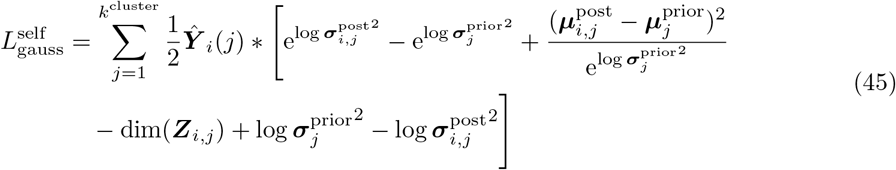

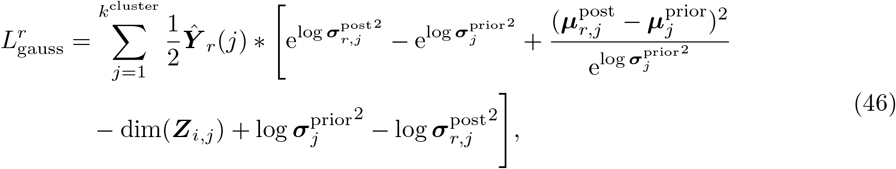

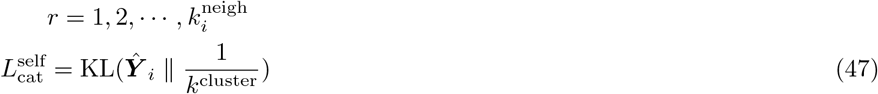

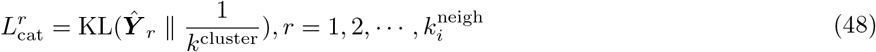

### Training strategies of stDyer

stDyer adopted a two-stage training process. In the first stage (1-50 epochs), stDyer was trained based on the spatial KNN graph and the Gaussian losses (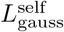 and 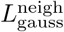) was ignored to enable the model to focus more on the reconstruction losses. In the second stage trained on dynamic graphs, we applied mclust [29] to group unitembeddings after the 50th epoch and generate pseudo spatial domain labels 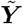 forall units. We then initialized predicted spatial domain labels ***Ŷ***using 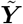using cross-entropy loss (Supplementary Equation (S3)) for 10 epochs (Supplementary Note S7). From the 61st to the 200th epoch, *L*_GMVAE_ was employed for model training and spatial domain clustering. We utilized an Adam optimizer to optimize these objective functions with a learning rate of 0.0001 and training epochs of 200. The prior distribution of unit domain labels was initialized as a uniform distribution and updated based on unit temporary spatial domain labels from the last epoch with a learning rate of 0.1.

### Spatial domain label refinement

To refine the spatial domains with outlier labels, we used sklearn.neural network.MLPClassifier to construct the autoencoder with mean squared error (MSE) as the loss function. For model training, the input and output of this autoencoder are spatial coordinates after min-max normalization and spatial domain labels from the modified GMVAE. We up-sampled the units from the smallest domain during model training to avoid this domain disappearing after refinement. This was done by scaling them up to the unit number of the second-smallest domains.

### Integrated gradient analysis

We incorporated the integrated gradients analysis [30]) in stDyer to identify SVGs. The IG vector was used to represent the effect of gene *g* in predicting the spatial domain label of unit *i*.

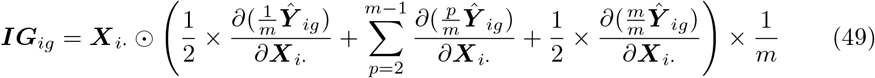

where *m* (50 by default) is the number of steps needed in the summation procedure in Equation (49) to estimate the definite integral. ***X*** and ***Ŷ*** are gene expression profiles of units and their predicted spatial domain labels, respectively. We averaged IG vectors of gene *g* across all units for each spatial domain to estimate its impact in prediction. Thus, SVGs for each domain were selected if their IGs within the top 50 largest IGs.

### Evaluation metric

We used Adjusted Rand Index (ARI) to evaluate the performance of spatial domain clustering [31]. ARI is a score computed between ground truth domain labels and predicted domain labels, ranging from -1 to 1. A higher ARI score indicates better clustering performance. ARI is defined as the following given the ground truth spatial domain ***A*** = (*a*_1_, *a*_2_, …, *a*_*t*_) and the predicted spatial domain ***B*** = (*b*_1_, *b*_2_, …, *b*_*p*_):

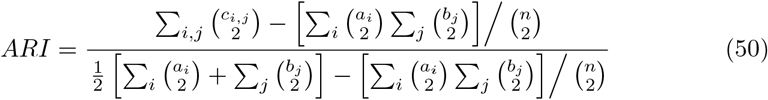

where *c*_*i*,*j*_ = *a*_*i*_ *b*_*j*_, and *n* is the number of units. We computed the p-value of ARIs on the multi-slice DLPFC dataset using the Wilcoxon rank-sum test with stat compare means function in ggpubr package [32]. The settings of hyper-parameters of stDyer and the other involved methods were shown in Supplementary Note S7 and Note S9.

## Data availability

The processed datasets below together with reproducible Jupyter notebooks are available at zenodo https://doi.org/10.5281/zenodo.11315102. The human dorsolateral prefrontal cortex dataset [1] is available in https://research.libd.org/spatialLIBD/ in h5 format. The osmFISH dataset [10] can be downloaded at http://linnarssonlab.org/osmFISH/availability/ in loom format. The STARmap dataset [21] is accessible at https://figshare.com/articles/dataset/STARmapdatasets/22565209. The zebrafish tumor dataset [11] is available at the Gene Expression Omnibus (GEO) under accession number ”GSE159709”. The Stereo-seq datasets [2] are downloaded from https://db.cngb.org/stomics/mosta/download/.

## Code availability

The source code of stDyer is publicly available at Github: https://github.com/ericcombiolab/stDyer.

## Authors’ contributions

L.Z conceived and supervised the project. K.X. and L.Z designed the method, conducted the experiments and wrote the manuscript. All authors reviewed and approved the manuscript.

## Supplementary information

There is a single supplementary file containing all supplementary materials.

## References

[1] Maynard, K.R., Collado-Torres, L., Weber, L.M., Uytingco, C., Barry, B.K., Williams, S.R., Catallini, J.L., Tran, M.N., Besich, Z., Tippani, M., Chew, J., Yin, Y., Kleinman, J.E., Hyde, T.M., Rao, N., Hicks, S.C., Martinowich, K., Jaffe, A.E.: Transcriptome-scale spatial gene expression in the human dorsolateral prefrontal cortex. Nature Neuroscience 24(3), 425–436 (2021) 10.1038/s41593-020-00787-0

[2] Chen, A., Liao, S., Cheng, M., Ma, K., Wu, L., Lai, Y., Qiu, X., Yang, J., Xu, J., Hao, S., Wang, X., Lu, H., Chen, X., Liu, X., Huang, X., Li, Z., Hong, Y., Jiang, Y., Peng, J., Liu, S., Shen, M., Liu, C., Li, Q., Yuan, Y., Wei, X., Zheng, H., Feng, W., Wang, Z., Liu, Y., Wang, Z., Yang, Y., Xiang, H., Han, L., Qin, B., Guo, P., Lai, G., Muñoz-Cánoves, P., Maxwell, P.H., Thiery, J.P., Wu, Q.-F., Zhao, F., Chen, B., Li, M., Dai, X., Wang, S., Kuang, H., Hui, J., Wang, L., Fei, J.-F., Wang, O., Wei, X., Lu, H., Wang, B., Liu, S., Gu, Y., Ni, M., Zhang, W., Mu, F., Yin, Y., Yang, H., Lisby, M., Cornall, R.J., Mulder, J., Uhlén, M., Esteban, M.A., Li, Y., Liu, L., Xu, X., Wang, J.: Spatiotemporal transcriptomic atlas of mouse organogenesis using dna nanoball-patterned arrays. Cell 0(0) (2022) 10.1016/j.cell.2022.04.003

[3] Zhu, Q., Shah, S., Dries, R., Cai, L., Yuan, G.-C.: Identification of spatially associated subpopulations by combining scrnaseq and sequential fluorescence in situ hybridization data. Nature Biotechnology 36(12), 1183–1190 (2018) 10.1038/nbt.4260

[4] Zhao, E., Stone, M.R., Ren, X., Guenthoer, J., Smythe, K.S., Pulliam, T., Williams, S.R., Uytingco, C.R., Taylor, S.E.B., Nghiem, P., Bielas, J.H., Gottardo, R.: Spatial transcriptomics at subspot resolution with bayesspace. Nature Biotechnology, 1–10 (2021) 10.1038/s41587-021-00935-2

[5] Li, Z., Zhou, X.: Bass: multi-scale and multi-sample analysis enables accurate cell type clustering and spatial domain detection in spatial transcriptomic studies. Genome Biology 23(1), 168 (2022) 10.1186/s13059-022-02734-7

[6] Thomas N. Kipf, Max Welling: Semi-supervised classification with graph convolutional networks. International Conference on Learning Representations (2016)

[7] Hu, J., Li, X., Coleman, K., Schroeder, A., Ma, N., Irwin, D.J., Lee, E.B., Shinohara, R.T., Li, M.: Spagcn: Integrating gene expression, spatial location and histology to identify spatial domains and spatially variable genes by graph convolutional network. Nature Methods, 1–10 (2021) 10.1038/s41592-021-01255-8

[8] Dong, K., Zhang, S.: Deciphering spatial domains from spatially resolved transcriptomics with an adaptive graph attention auto-encoder. Nature Communications 13(1), 1739 (2022) 10.1038/s41467-022-29439-6

[9] Long, Y., Ang, K.S., Li, M., Chong, K.L.K., Sethi, R., Zhong, C., Xu, H., Ong, Z., Sachaphibulkij, K., Chen, A., Zeng, L., Fu, H., Wu, M., Lim, L.H.K., Liu, L., Chen, J.: Spatially informed clustering, integration, and deconvolution of spatial transcriptomics with graphst. Nature Communications 14(1), 1155 (2023) 10.1038/s41467-023-36796-3

[10] Codeluppi, S., Borm, L.E., Zeisel, A., La Manno, G., van Lunteren, J.A., Svensson, C.I., Linnarsson, S.: Spatial organization of the somatosensory cortex revealed by osmfish. Nature Methods 15(11), 932–935 (2018) 10.1038/s41592-018-0175-z

[11] Hunter, M.V., Moncada, R., Weiss, J.M., Yanai, I., White, R.M.: Spatially resolved transcriptomics reveals the architecture of the tumor-microenvironment interface. Nature Communications 12(1), 6278 (2021) 10.1038/s41467-021-26614-z

[12] Cang, Z., Ning, X., Nie, A., Xu, M., Zhang, J.: Scan-it: Domain segmentation of spatial transcriptomics images by graph neural network. BMVC: proceedings of the British Machine Vision Conference. British Machine Vision Conference 32 (2021)

[13] Ren, H., Walker, B.L., Cang, Z., Nie, Q.: Identifying multicellular spatiotemporal organization of cells with spaceflow. Nature Communications 13(1), 4076 (2022) 10.1038/s41467-022-31739-w

[14] Xu, C., Jin, X., Wei, S., Wang, P., Luo, M., Xu, Z., Yang, W., Cai, Y., Xiao, L., Lin, X., Liu, H., Cheng, R., Pang, F., Chen, R., Su, X., Hu, Y., Wang, G., Jiang, Q.: Deepst: identifying spatial domains in spatial transcriptomics by deep learning. Nucleic Acids Research 50(22), 131 (2022) 10.1093/nar/gkac901

[15] Yuan, Z., Zhao, F., Lin, S., Zhao, Y., Yao, J., Cui, Y., Zhang, X.-Y., Zhao, Y.: Benchmarking spatial clustering methods with spatially resolved transcriptomics data. Nature Methods, 1–11 (2024) 10.1038/s41592-024-02215-8

[16] Dilokthanakul, N., Mediano, P.A.M., Garnelo, M., Lee, M.C.H., Salimbeni, H., Arulkumaran, K., Shanahan, M.: Deep Unsupervised Clustering with Gaussian Mixture Variational Autoencoders. https://arxiv.org/pdf/1611.02648

[17] Jiang, Z., Zheng, Y., Tan, H., Tang, B., Zhou, H.: Variational deep embedding: An unsupervised and generative approach to clustering. In: Proceedings of the Twenty-Sixth International Joint Conference on Artificial Intelligence, IJCAI-17, pp. 1965–1972 (2017). 10.24963/ijcai.2017/273

[18] Yang, L., Cheung, N.-M., Li, J., Fang, J.: Deep clustering by gaussian mixture variational autoencoders with graph embedding. In: 2019 International Conference on Computer Vision, pp. 6439–6448. IEEE, Piscataway, NJ (2019). 10.1109/ICCV.2019.00654. https://openaccess.thecvf.com/content_ICCV_2019/papers/Yang_Deep_Clustering_by_Gaussian_Mixture_Variational_Autoencoders_With_Graph_Embedding_ICCV_2019_paper.pdf Accessed 2021/6/23

[19] Petar Veličković, Guillem Cucurull, Arantxa Casanova, Adriana Romero, Pietro Liò, Yoshua Bengio: Graph attention networks. International Conference on Learning Representations (2022)

[20] Rebecca, V.W., Nicastri, M.C., Fennelly, C., Chude, C.I., Barber-Rotenberg, J.S., Ronghe, A., McAfee, Q., McLaughlin, N.P., Zhang, G., Goldman, A.R., Ojha, R., Piao, S., Noguera-Ortega, E., Martorella, A., Alicea, G.M., Lee, J.J., Schuchter, L.M., Xu, X., Herlyn, M., Marmorstein, R., Gimotty, P.A., Speicher, D.W., Winkler, J.D., Amaravadi, R.K.: Ppt1 promotes tumor growth and is the molecular target of chloroquine derivatives in cancer. Cancer Discovery 9(2), 220–229 (2019) 10.1158/2159-8290.CD-18-0706

[21] Wang, X., Allen, W.E., Wright, M.A., Sylwestrak, E.L., Samusik, N., Vesuna, S., Evans, K., Liu, C., Ramakrishnan, C., Liu, J., Nolan, G.P., Bava, F.-A., Deisseroth, K.: Three-dimensional intact-tissue sequencing of single-cell transcriptional states. Science 361(6400) (2018) 10.1126/science.aat5691

[22] Baldarelli, R.M., Smith, C.L., Ringwald, M., Richardson, J.E., Bult, C.J.: Mouse genome informatics: an integrated knowledgebase system for the laboratory mouse. Genetics 227(1) (2024) 10.1093/genetics/iyae031

[23] Safran, M., Rosen, N., Twik, M., BarShir, R., Stein, T.I., Dahary, D., Fishilevich, S., Lancet, D.: The genecards suite. In: Practical Guide to Life Science Databases, pp. 27–56. Springer, ??? (uuuu-uuuu). 10.1007/978-981-16-5812-92. https://link.springer.com/chapter/10.1007/978-981-16-5812-9 2

[24] Tutukova, S., Tarabykin, V., Hernandez-Miranda, L.R.: The role of neurod genes in brain development, function, and disease. Frontiers in Molecular Neuroscience 14, 662774 (2021) 10.3389/fnmol.2021.662774

[25] Yan, W., Long, P., Chen, T., Liu, W., Yao, L., Ren, Z., Li, X., Wang, J., Xue, J., Tao, Y., Zhang, L., Zhang, Z.: A natural occurring mouse model with adgrv1 mutation of usher syndrome 2c and characterization of its recombinant inbred strains. Cellular physiology and biochemistry: international journal of experimental cellular physiology, biochemistry, and pharmacology 47(5), 1883|1897 (2018) 10.1159/000491068

[26] He, K., Zhang, X., Ren, S., Sun, J.: Deep residual learning for image recognition, pp. 770–778 (2016). https://openaccess.thecvf.com/content_cvpr_2016/html/He_Deep_Residual_Learning_CVPR_2016_paper.html

[27] Wolf, F.A., Angerer, P., Theis, F.J.: Scanpy: large-scale single-cell gene expression data analysis. Genome Biology 19(1), 15 (2018) 10.1186/s13059-017-1382-0

[28] Liu, X., Zeira, R., Raphael, B.J.: Partial alignment of multislice spatially resolved transcriptomics data. Genome Research 33(7), 1124–1132 (2023) 10.1101/gr.277670.123

[29] Scrucca, L., Fop, M., Murphy, T.B., Raftery, A.E.: mclust 5: Clustering, classification and density estimation using gaussian finite mixture models. The R journal 8(1), 289–317 (2016)

[30] Sundararajan, M., Taly, A., Yan, Q.: Axiomatic attribution for deep networks. In: Precup, D., Teh, Y.W. (eds.) Proceedings of the 34th International Conference on Machine Learning. Proceedings of Machine Learning Research, vol. 70, pp. 3319–3328. PMLR, New York, New York, USA (2017). https://proceedings.mlr.press/v70/sundararajan17a.html

[31] Hubert, L., Arabie, P.: Comparing partitions. Journal of Classification 2(1), 193–218 (1985) 10.1007/BF01908075

[32] Kassambara, A.: Ggpubr: ‘ggplot2’ Based Publication Ready Plots. (2023). R package version 0.6.0. https://rpkgs.datanovia.com/ggpubr/

[33] Dries, R., Zhu, Q., Dong, R., Eng, C.-H.L., Li, H., Liu, K., Fu, Y., Zhao, T., Sarkar, A., Bao, F., George, R.E., Pierson, N., Cai, L., Yuan, G.-C.: Giotto: a toolbox for integrative analysis and visualization of spatial expression data. Genome Biology 22(1), 78 (2021) 10.1186/s13059-021-02286-2

