## Supplementary for "stDyer enables spatial domain clustering with dynamic graph embedding"

### Supplementary Information

#### S1 Spatial domain clustering results

##### S1.1 DLPFC dataset

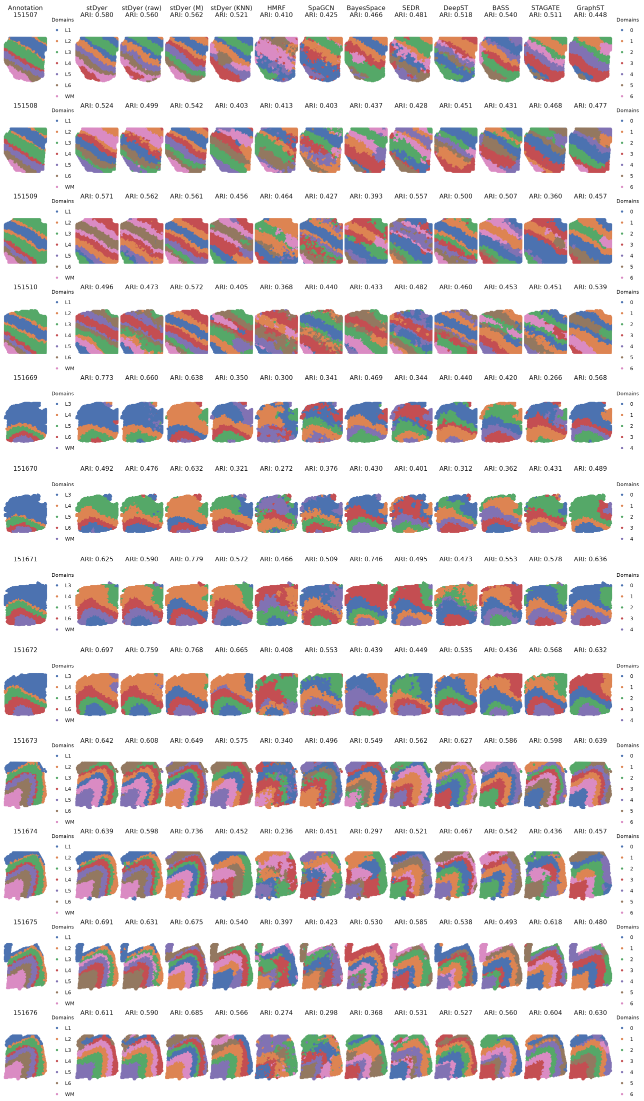


**Supplementary Figure S1**: The annotation and clustering result visualization and ARI score of each method. ARI: adjusted rand index.

##### S1.2 Stereo-seq dataset

**Supplementary Table S1**: ARI results for Stereo-seq dataset.

| Slice ID | Unit number | stDyer | SpaGCN | BayesSpace | SEDR | GraphST | BASS | DeepST | STAGATE |
|---|---|---|---|---|---|---|---|---|---|
| 1 | 59119 | 0.381287 | 0.361367 | 0.371171 | 0.343704 | 0.359263 | NA | NA | NA |
| 2 | 73883 | 0.376097 | 0.307762 | 0.291434 | 0.325156 | 0.346139 | NA | NA | NA |
| 13 | 77688 | 0.356095 | 0.330281 | 0.309785 | 0.288363 | 0.312356 | NA | NA | NA |
| 12 | 94665 | 0.362656 | 0.351810 | 0.336085 | 0.328799 | 0.299853 | NA | NA | NA |
| 4 | 99200 | 0.515413 | 0.425840 | 0.386995 | 0.432986 | NA | NA | NA | NA |
| 11 | 109383 | 0.421177 | 0.374813 | 0.395944 | 0.329744 | NA | NA | NA | NA |
| 3 | 110460 | 0.410080 | 0.361773 | 0.335751 | 0.396288 | NA | NA | NA | NA |
| 10 | 113947 | 0.416465 | 0.385074 | 0.385576 | 0.374271 | NA | NA | NA | NA |
| 8 | 120815 | 0.409339 | 0.368711 | 0.330293 | 0.347299 | NA | NA | NA | NA |
| 9 | 129423 | 0.389422 | 0.365286 | 0.379837 | 0.299194 | NA | NA | NA | NA |
| 5 | 132455 | 0.394218 | 0.376458 | 0.382694 | NA | NA | NA | NA | NA |
| 6 | 137113 | 0.315178 | 0.306262 | 0.303599 | NA | NA | NA | NA | NA |
| 7 | 155047 | 0.358662 | 0.254089 | 0.284076 | NA | NA | NA | NA | NA |

#### S2 Spatially variable gene visualization

The top 5 genes with the largest integrated gradients of each predicted domain were visualized.

#### S2.1 DLPFC dataset

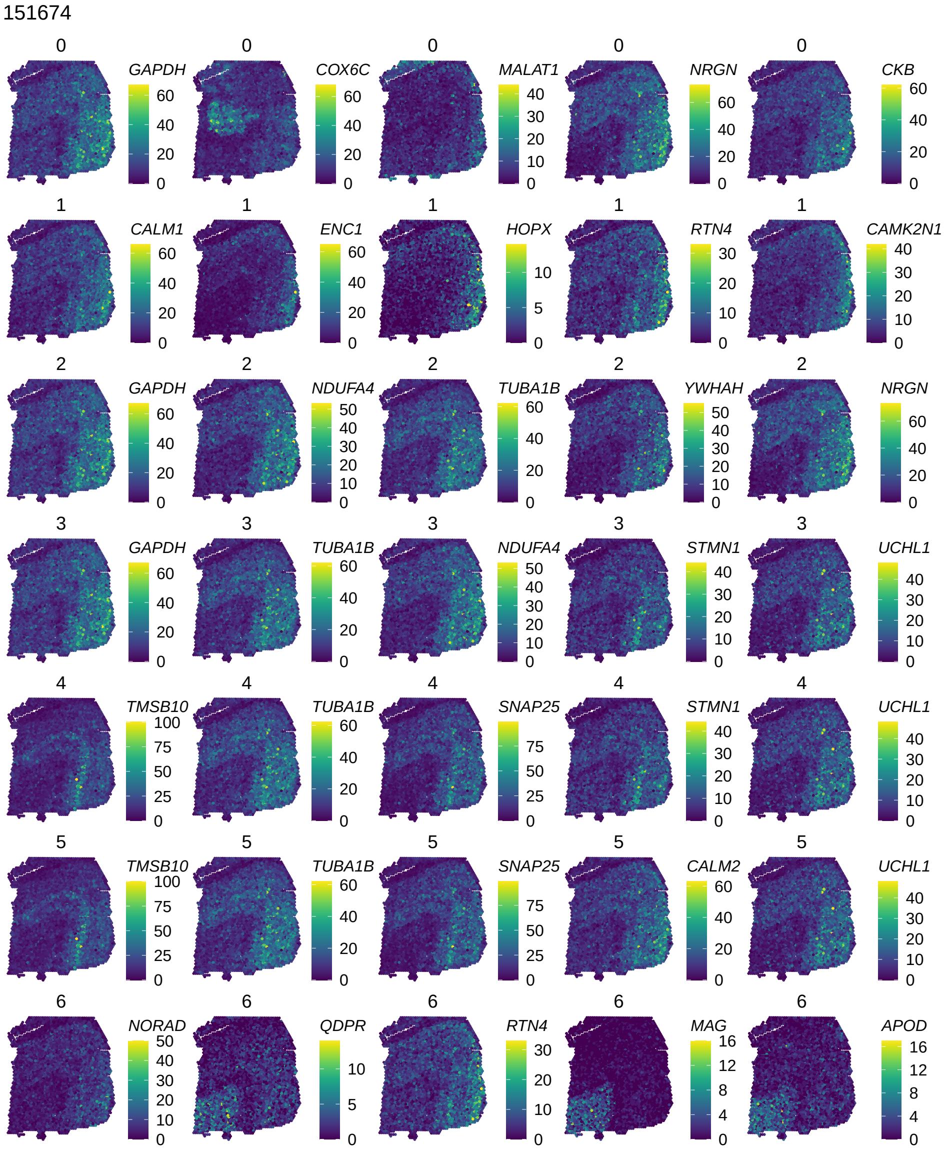


**Supplementary Figure S2**: The top 5 SVGs of slice 151674 on each predicted domain. SVG, spatially variable gene.

| Domain | Spatially variable genes |
|---|---|
| 0 | *GAPDH, COX6C, MALAT1, NRGN, CKB, CAMK2N1, GNAS, SLC1A2, NDUFA4, HSP90AB1, CALM3, ALB, SNAP25, ACTB, ENO2, EEF1A2, BASP1, AP2M1, PRKAR1B, MYL9, GNG3, MEG3, SNCG, ATP1A3, SLC17A7, WSB2, S100G, YWHAB, TMSB10, PDE11A, TAGLN3, CALM1, ADAMTS1, COL3A1, VAMP2, BRAF, STMN3, ACTA2, NDST3, NSG2, NPTXR, MGP, GABRA1, ZNF350-AS1, VSTM2A, YWHAZ, OLFM1, ATP1B1, COL6A1, FAM49A* |
| 1 | *CALM1, ENC1, HOPX, RTN4, CAMK2N1, ARPP19, YWHAH, NDFIP1, PCDH8, HPCAL1, PTGDS, ATP1A1, NECAB1, CKB, CALY, SOWAHA, FKBP1A, CCK, NAT8L, STXBP1, VSNL1, SERPINE2, ATP2B2, CAMK2A, TSC22D1, SLC6A1, SPARCL1, BAIAP3, NRGN, LAMP5, CYP46A1, DOCK4, PHYHIP, PPFIA2, GRIA2, GRIN1, RASGRF2, SARAF, PRKCA, FABP4, GRIA4, CUX2, CNKSR2, YJEFN3, CALB1, R3HDM1, HAPLN4, NRXN1, RBP4, DPP6* |
| 2 | *GAPDH, NDUFA4, TUBA1B, YWHAH, NRGN, ACTB, NEFM, STMN1, ATP1B1, CKB, UCHL1, NORAD, RTN1, CCK, ATP1A3, NEFL, ATP5F1B, GAP43, SNCB, MEF2C, MDH1, CPLX1, TUBB2A, CLTC, CRYM, YWHAG, STXBP1, BASP1, GPX3, CALM2, MAP1B, SNRPN, SLC12A5, RTN3, ENO2, TSC22D1, HOPX, GNAS, LMO4, SNCG, NDRG4, OXR1, RAB3A, NUDT4, DCLK1, CABP1, EEF1A2, DNM1, SPTBN1, STMN2* |
| 3 | *GAPDH, TUBA1B, NDUFA4, STMN1, UCHL1, ACTB, NORAD, MAP1B, ATP1B1, NEFL, RTN1, NRGN, NEFM, SNCB, RTN3, YWHAH, SNRPN, STXBP1, YWHAG, SNAP25, MEF2C, CLTC, ATP1A1, GPX3, STMN2, NDRG4, CCK, SARAF, CPLX1, HSPA8, VAMP1, ATP1A3, PGK1, ATP5F1B, INA, VSNL1, BASP1, ENO2, LMO4, DCLK1, OXR1, SNCG, TSC22D1, CRYM, BEX5, MDH1, PTGDS, NDFIP1, CABP1, NUDT4* |
| 4 | *TMSB10, TUBA1B, SNAP25, STMN1, UCHL1, YWHAG, MAP1B, ACTB, RTN3, TUBB2A, GNAS, NDUFA4, SYT1, CALM2, NDRG4, HSPA8, SNCB, DIRAS2, PCP4, GAPDH, STMN2, BASP1, SNCG, NRN1, NEFL, VSNL1, SNRPN, CPLX1, PRKAR1B, ATP1B1, PFKP, MDH1, NEFM, ATP5MD, TAGLN3, CALM3, NAP1L5, STMN3, AP2M1, SPTBN1, TUBA4A, TSPAN7, ANXA6, NUDT4, STXBP1, SCGB2A2, NAP1L3, ATP5F1B, TTC9B, GHITM* |
| 5 | *TMSB10, TUBA1B, SNAP25, CALM2, UCHL1, SYT1, GNAS, STMN1, YWHAG, COX6C, DIRAS2, DKK3, CALM3, GPM6A, CHN1, SCGB1D2, SLC17A7, CCK, SNCB, RTN3, TSPAN7, RTN1, SCGB2A2, BASP1, MBP, HSP90AB1, PRKAR1B, STMN3, KRT19, TUBB2A, GABRA5, SV2B, KRT17, THY1, SARAF, MAP1B, CPB1, PPP3CA, STMN2, NEFL, NRGN, VSNL1, SLC24A2, CNP, NCALD, NELL2, ATP2B1, KLC1, AK5, MOAP1* |
| 6 | *NORAD, QDPR, RTN4, MAG, APOD, QKI, ERMN, SELENOP, RNASE1, MARCKSL1, LHPP, MOBP, SCD, SLC44A1, FA2H, SIRT2, CLDND1, MAP1B, ABCA2, DNER, SOX10, PLP1, HSPA2, RTN1, TTYH2, MAL, PPP1R14A, AMER2, SEPT4, PLPPR1, PLLP, SPP1, PREX1, MBP, PSEN1, RNF13, CA2, TSC22D4, GPM6B, GLTP, DAAM2, FEZ1, PTGDS, GFAP, PIP4K2A, CERCAM, AQP1, MORF4L2, TSPYL1, OPALIN* |

**Supplementary Table S2**: The top 50 SVGs of slice 151674 on each predicted domain.

#### S2.2 STARmap dataset

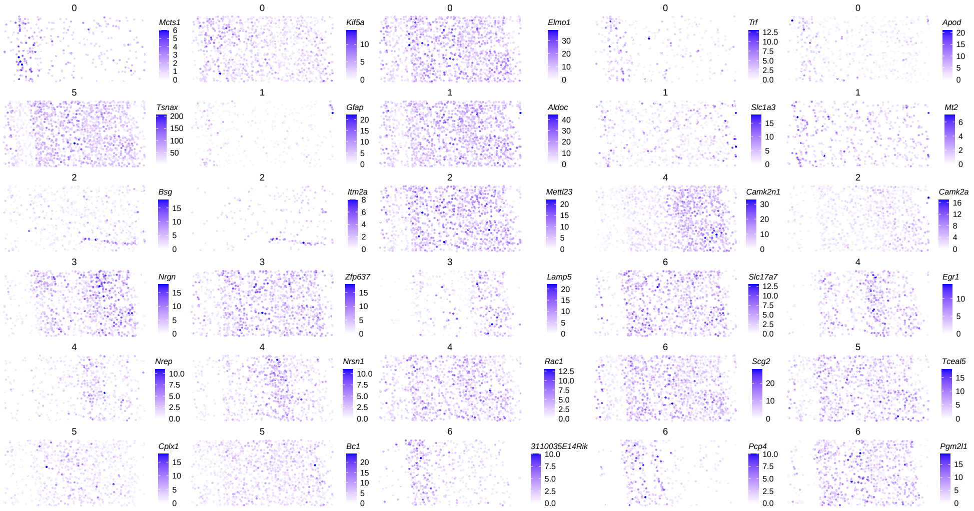


**Supplementary Figure S3**: The top 5 SVGs of STARmap dataset on each predicted domain. SVG, spatially variable gene.

| Domain | Spatially variable genes |
|---|---|
| 0 | *Mcts1, Kif5a, Elmo1, Trf, Apod, Gatm, Plp1, Gstm6, Sept4, Mal, Gsn, Mbp, Ptgds, Syt6, Trpv6, Hmgcs1, Fam19a1, Bc1, Cd9, Dusp6, Stmn4, Nudt4, Plekhb1, Cmtm5, Fbxl3, Fbxo44, Qk, Eid1, Tmem176a, Cend1, Sirt2, Sv2b, Cryab, Teddm3, Rbm18, Slc24a2, Synpr, Fto, Nek7, Zfp637, Neurod2, Adarb2, Bcl6, Hlf, Pcdha12, Glul, Pllp, Ccdc80, Rasgrp1, Myl4* |
| 1 | *Tsnax, Gfap, Aldoc, Slc1a3, Mt2, Mt1, Gstm1, Atp1a2, Apoe, Sdc4, Pantr1, Ppp1r3c, Htra1, Slc1a2, Pon2, Dbi, Oxt, Cd63, Mettl23, Pcdhga3, Aamp, Sox9, Efemp1, Mia, S100a10, Ddit3, Ndrg2, Sparcl1, Cldn10, Mfge8, Gpr88, Bcan, S100a1, Cpne2, Dtwd1, Gstm5, Timp4, Avp, Slc25a36, Ifi27, Acsbg1, Rbp4, Aqp4, Gpr37l1, Myh1, Bsg, Ifit3, Fabp7, Agrp, Mrpl33* |
| 2 | *Bsg, Itm2a, Mettl23, Camk2n1, Camk2a, Fosb, Ifitm2, Dusp6, Hlf, Slc2a13, Tsnax, Glul, Sepp1, Zdhhc24, Hpcal4, Ly6e, B2m, Dusp1, Mrpl22, Bcl6, Fam163b, Aebp1, Itih5, Nrg1, Sparc, Gm2a, Scg2, Nr2f2, Tmbim6, Clic1, Sdcbp, Hs3st1, Fkbp1a, S100a16, Grm3, Ndrg2, Tmem176a, Gpx3, Dkk3, Sema3c, Gm15881, Pcdhga3, Sostdc1, Cldn5, Gpx1, Pcdhga5, Pcdhb10, Thrsp, Btg2, Mdga1* |
| 3 | *Nrgn, Zfp637, Camk2n1, Lamp5, Slc17a7, Enc1, Elmo1, Ncdn, Hpcal4, Egr3, 2900055J20Rik, Gucy1b3, Mat2b, Pcsk2, Gria3, Hhatl, Cbln2, Atraid, Cpne5, Fxyd7, Hdac9, Atp2b4, Homer1, Ddit4l, Pgm2l1, Pcdhb9, Pdzrn3, Efna5, Hpca, Fkbp1a, Tpm1, Nudt2, Cyb561, Chodl, Cnr1, Cpne9, Osbpl1a, Agtr1a, Map2k1, Cd34, Dusp1, Rbfox3, Tsnax, Calb1, Pip5k1b, Exosc9, 1700086L19Rik, Fam19a2, Pea15a, Pcdhb6* |
| 4 | *Camk2n1, Egr1, Nrep, Nrsn1, Rac1, Nrgn, Zfp637, Arc, Zmat4, Slc17a7, Scg2, Snca, Nrn1, Pdp1, Fbxw11, Whrn, Egr3, Fos, Mef2c, Cyb561, Ncdn, Sdcbp, Btbd3, Pgm2l1, Rasgrp1, Stx1a, Mettl23, Cadps2, Nudt4, Rcan2, Elmo1, Egr2, Satb2, Camk2a, Sat1, Ppp3cb, Mtfp1, Epha4, Bc1, Ras-grf1, 2900055J20Rik, Car10, Syn1, Nell1, Rbfox3, Atp9a, 2900092D14Rik, Map2k1, Herc3, Nipsnap3b* |
| 5 | *Tsnax, Scg2, Tceal5, Cplx1, Bc1, Htra1, Efr3a, Ndufs2, Rab3c, St6galnac5, Snurf, Pcp4, Pgm2l1, Itm2c, Ogfrl1, Eid1, Luzp2, Slc17a7, Gad1, Pea15a, Aldoc, Atp1a2, Nov, Kcnc2, Tox, Gpr22, 3110035E14Rik, Trim32, Pcdhga3, Mal2, Plcxd3, Epha7, Cd9, Snx10, Pcsk2, Lgmn, Slc50a1, Msmo1, Otof, Tmsb10, Lmo4, Snca, Gda, Elp3, Lman1l, Cadps2, Crh, Med21, Egln3, Rit2* |
| 6 | *Scg2, Slc17a7, 3110035E14Rik, Pcp4, Pgm2l1, Arc, Ogfrl1, Rac1, Nrgn, Zfp637, Snca, Hpcal4, Egr1, Syn1, Rasgrp1, Tle4, Plcxd3, Garnl3, Arpp19, Tceal5, Prkcg, Pcsk2, Fxyd7, Rprm, Arhgap25, Rap1gds1, Bc1, Klf10, Cyb561, Lmo4, Stx1a, Ppp3cb, Col6a1, Mpi, Rcan2, Lmo3, St3gal5, Slc6a15, Homer1, Arx, Rgs2, Med10, Map2k1, Kifc2, Rbfox3, Egr3, Sv2b, Snurf, Ndufs2, Efhd2* |

**Supplementary Table S3**: The top 50 SVGs of STARmap dataset on each predicted domain.

#### S2.3 Zebrafish tumor dataset

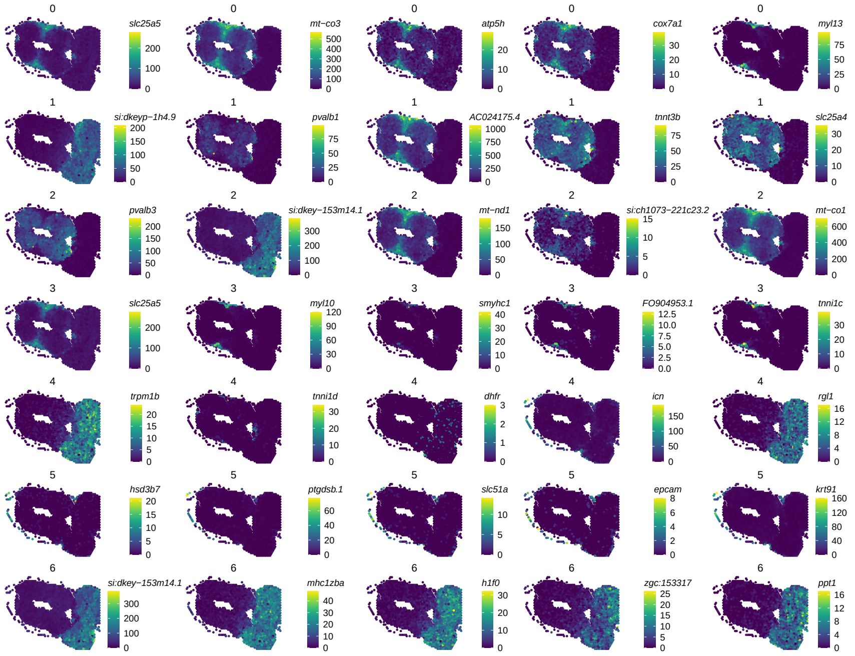


**Supplementary Figure S4**: The top 5 SVGs of zebrafish dataset on each predicted domain. SVG, spatially variable gene.

| Domain | Spatially variable genes |
|---|---|
| 0 | *slc25a5, mt-co3, atp5h, cox7a1, myl13, CABZ01076275.1, tnnc1b, mt-co2, myl10, arg2, mt-cyb, tpm2, rpl14, cox7a3, pvalb7, mb, mybpc2a, tnni1al, acadvl, tnnt2e, tcap, ACTC1 (1 of many), atp2a1, flnca, epas1b, atp2a2a, adck3, nme2b.2, pdk2b, ndufa4l, si:dkey-206m15.8, smyhc2, si:ch211-5k11.8, IDH2, mt-nd4l, mt-nd5, ppdpfa, tnni1c, mylk3, slc25a33, pdk2a, sorbs1, si:dkey-181m9.5, mt-nd2, prelid3b, nmrk2, zgc:114041, casq1b, mlip, hspb1* |
| 1 | *si:dkeyp-1h4.9, pvalb1, AC024175.4, tnnt3b, slc25a4, ckma, atp2a1l, si:ch73-367p23.2, rps18, zgc:152791, ttn.2, pvalb2, krt91, actb1, hmgn2, icn, actn3a, nme2b.2, jph2, CU984600.2, myhc4, slc14a2, mylpfa, ndufa4l, myhz1.1-1, ilf2, CH25H, pgam2, pvalb3, naca, mllt11, tuba8l4, actn3b, hsp90ab1, ehbp1l1a, dusp6, hunk, mt-atp6, fkbp10b, col1a1b, krt5, anxa2a, si:ch211-81a5.8, ucp3, ryr3, anxa1a, b2m, calm3b, mt-nd1, efr3bb* |
| 2 | *pvalb3, si:dkey-153m14.1, mt-nd1, si:ch1073-221c23.2, mt-co1, tnni2a.4, AC024175.17, atp1a2a, atp5g1, ckmt2a, myl1, tpma, chchd10, cox5b2, tnnc2, trim63a, ak1, IDH2, nln, chrne, pkmb, tnni2a.3, col1a1a, frmd8, mt-atp6, ba1-1, cpt1b, mylz3, mhc1zba, myhc4, pvalb2, slc25a4, egfra, LRRC2, zgc:158463, sod1, vwde, tubb4b, fam136a, sdc4, marcksl1b, lpin1, actn3b, slco2a1, si:dkey-188i13.9, abca1a, gulp1a, ywhabl, cacna2d1a, pgam2* |
| 3 | *slc25a5, myl10, smyhc1, FO904953.1, tnni1c, tnnt2e, si:dkey-206m15.8, tnnc1b, tpm2, atp2a2a, myl13, tnni1d, mybpc3, fstl1a, ankrd1a, nmrk2, fhl1a, mylk3, arg2, myom3, rbp7b, zgc:114041, il15l-1, CABZ01076275.1, hbaa1, atp5h, rnf207b, tnfrsf11a, syne2b, CU633479.1, rpl14, hhatlb, tnni1al, xirp1, pth1b, pank1a, si:dkey-71d15.2, gls2b, slc25a33, nr4a3, mt-nd3, acadvl, si:ch211-247n2.1, rcan3, eci2, map1lc3cl, dnajb6b, ech1, lipg, si:dkey-153m14.1* |
| 4 | *trpm1b, tnni1d, dhfr, icn, rgl1, wu:fi09b08-6, h2afva, prdx2, tuba8l, ebi3, si:rp71-45k5.4, perp, nop58, cbfb, apoda.1, zgc:163073, vmhc, krt91, paics, krt18, col1a1a, arap3, ppiab, anxa1a, cd74a, spp1, zgc:193505, hexb, pdlim5b, si:ch211-251b21.1, tktb, ing4, hnrnpa1b, rps18, ntrk3a, hmga1a, ywhaz, ifitm1, myo5b, emp3b, tomm20a, pelp1, IFI30 (1 of many), si:dkey-188i13.9, mrgbp, gpnmb, rnpep, zgc:172244, pnp4a, atp6v1aa* |
| 5 | *hsd3b7, ptgdsb.1, slc51a, epcam, krt91, tmem97, icn, krt5, hsd17b7, cyp8b2, anxa1a, apoeb, si:dkey-7c18.24, cfl1l, abcb5, perp, ptgdsb.2, aqp3a, si:dkey-193p11.2, zgc:111983, si:dkey-153m14.1, itga6b, cldnb, krt15, aldob, si:ch211-195b11.3, gig2l, itgb4, sult2st3, TMSB15A, samhd1, agr1, agr2, si:ch211-69b7.6, spaca4l, ebp, si:dkey-248g15.3, s100w, icn2, atp1a3b, msmo1, si:ch211-241e1.3, si:dkey-262k9.2, cd74b, cldna, si:zfos-1011f11.1, lgals3bpa, fdps, endouc, porb* |
| 6 | *si:dkey-153m14.1, mhc1zba, h1f0, zgc:153317, ppt1, zgc:101846, si:dkey-226k3.4, arrdc3b, sox5, zgc:158463, stat1b, cyfip1, snrpd1, pmela, ctps1a, ranbp1, eif5a2, SPAG9 (1 of many), actr1, stmn1a, ptprea, polr2k, tacc1, mxra8b, si:dkey-188i13.7, eno1a, pgam1a, znf395a, sub1a, LIMD1, ccnb1, hsp90aa1.2, lmnb2, h3f3a, ivns1abpa, nsun2, cdk1, phf5a, cirbpb, hcfc1a, zgc:171704, mdkb, psma3, dynll1, tfap2e, prkcbp1l, skiv2l2, atox1, eef1db, ighv1-4* |

**Supplementary Table S4**: The top 50 SVGs of zebrafish dataset on each predicted domain.

#### S3 Multi-slice grouping

**Supplementary Table S5**: Multi-slice grouping for DLPFC dataset.

| Group 1 | Group 2 | Group 3 |
|---|---|---|
| 151507 | 151669 | 151673 |
| 151508 | 151670 | 151674 |
| 151509 | 151671 | 151675 |
| 151510 | 151672 | 151676 |

### S4 Network structure

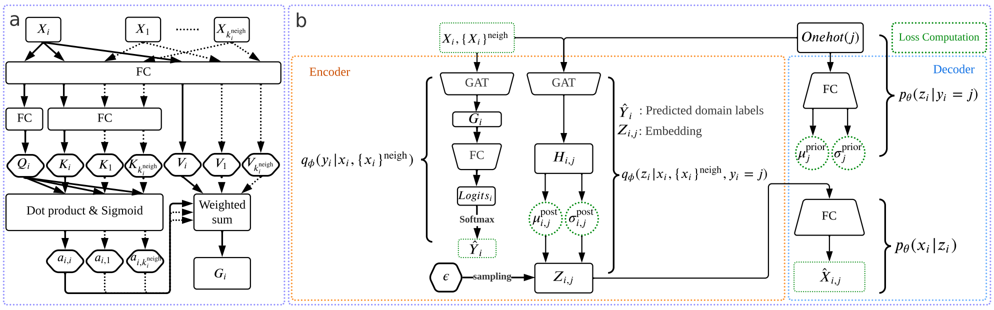


**Supplementary Figure S5**: (**a**) GAT structure. GAT first takes the target unit and its neighbors' expression profiles to generate query (Q), keys (K) and values (V) and uses the first two to obtain attention scores (a). The values are then weighted by the attention scores to generate the aggregated context. (**b**) Network structure. The backbone of stDyer is a Gaussian mixture variational autoencoder with detailed symbol definition in the Methods part. The encoder component integrates the graph attention mechanism to predict domain labels (left) and domain-specific embeddings (middle). The decoder component generates the prior Gaussian mixture parameters (top) and takes embeddings (middle) from the encoder to reconstruct the input (bottom). GAT, graph attention network.

### S5 Graph construction

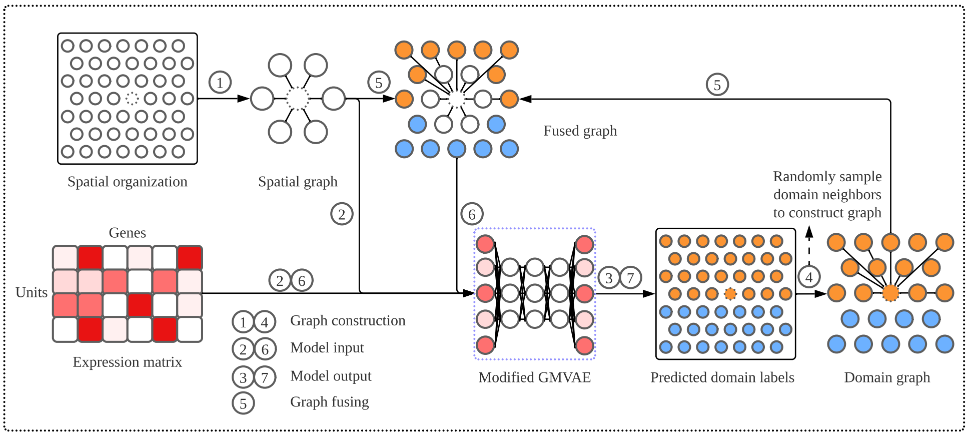


**Supplementary Figure S6**: Graph construction procedures. The circled numbers indicate the order of the data flow. Graph is first constructed by KNN and domain labels can be predicted to obtain domain neighbors. Domain neighbors can then be fused into the KNN graph to optimize the input graph iteratively.

### S6 Proof of lower bound

$\mathrm{JS}_{\theta,\phi}^{\mathrm{smooth}(r)}$ was the lower bound of $G_{\theta,\phi}^{r}$ (Equation (S2)) as:

$$G_{\theta,\phi} = \frac{1}{k_i^{\mathrm{neigh}}} \sum_{r=1}^{k_i^{\mathrm{neigh}}} G_{\theta,\phi}^{r} \tag{S1}$$

$$\begin{aligned}
G_{\theta,\phi}^{r} &= \frac{1}{2}\Bigg( E_{q_\phi(y_i,\boldsymbol{z}_i|\boldsymbol{x}_i,\{\boldsymbol{x}_i\}^{\mathrm{neigh}})} \log \frac{q_\phi(y_i,\boldsymbol{z}_i|\boldsymbol{x}_i,\{\boldsymbol{x}_i\}^{\mathrm{neigh}})}{p_\theta(y_i,\boldsymbol{z}_i|\boldsymbol{x}_i)} \\
&\quad + E_{q_\phi(y_r,\boldsymbol{z}_r|\boldsymbol{x}_r,\{\boldsymbol{x}_r\}^{\mathrm{neigh}})} \log \frac{q_\phi(y_r,\boldsymbol{z}_r|\boldsymbol{x}_r,\{\boldsymbol{x}_r\}^{\mathrm{neigh}})}{p_\theta(y_i,\boldsymbol{z}_i|\boldsymbol{x}_i)} \Bigg) \\
&= \frac{1}{2}\Bigg( E_{q_\phi(y_i,\boldsymbol{z}_i|\boldsymbol{x}_i,\{\boldsymbol{x}_i\}^{\mathrm{neigh}})} \log \frac{q_\phi(y_i,\boldsymbol{z}_i|\boldsymbol{x}_i,\{\boldsymbol{x}_i\}^{\mathrm{neigh}})}{M} \\
&\quad + E_{q_\phi(y_r,\boldsymbol{z}_r|\boldsymbol{x}_r,\{\boldsymbol{x}_r\}^{\mathrm{neigh}})} \log \frac{q_\phi(y_r,\boldsymbol{z}_r|\boldsymbol{x}_r,\{\boldsymbol{x}_r\}^{\mathrm{neigh}})}{M} \Bigg) + E_M \log \frac{M}{p_\theta(y_i,\boldsymbol{z}_i|\boldsymbol{x}_i)} \\
&= \mathrm{JS}_{\theta,\phi}^{\mathrm{smooth}(r)} + \mathrm{KL}(M \parallel p_\theta(y_i,\boldsymbol{z}_i|\boldsymbol{x}_i)) \\
&\geq \mathrm{JS}_{\theta,\phi}^{\mathrm{smooth}(r)}
\end{aligned} \tag{S2}$$

where $M = \frac{1}{2}[q_\phi(y_i, \boldsymbol{z}_i|\boldsymbol{x}_i, \{\boldsymbol{x}_i\}^{\text{neigh}}) + q_\phi(y_r, \boldsymbol{z}_r|\boldsymbol{x}_r, \{\boldsymbol{x}_r\}^{\text{neigh}})]$.

#### S7 Hyper-parameter settings

We set the learning rate to 0.001 empirically for faster convergence and stability. A higher learning rate like 0.01 could result in numerical instabilities while a lower learning rate below 0.0001 significantly reduced the learning efficiency. The batch size was set to 1,024 by default for faster convergence and training speed. Notably, the batch size could be adjusted to match the available graphics memory such that large datasets could also be run on mid-end consumer-level graphics cards with limited graphics memory.

We trained stDyer with 3 stages for 50, 10, and 140 epochs, respectively. In the first stage, we observed that 50 epochs were enough to minimize reconstruction loss by a large margin for learning embedding that could reconstruct the input well. For the second stage, we used 10 epochs with 1.01 as the threshold in the indicator function of the cross-entropy loss part to learn the pseudo labels of mclust for initializing the domain label predictor of stDyer. A higher threshold can stop stDyer from optimizing domain labels due to overfitting while a lower threshold can require more epochs for this stage to initialize stDyer well. For the third stage, we used the remaining 140 epochs out of 200 epochs in total to optimize the network to obtain the final predicted spatial domains.

$$\text{CE}(\hat{\boldsymbol{Y}}_i, \tilde{\boldsymbol{Y}}_i) = -\sum_{j=1}^{k^{\text{cluster}}} \tilde{\boldsymbol{Y}}_i(j) \log \hat{\boldsymbol{Y}}_i(j) \tag{S3}$$

$$I_{\text{CE}}^i = \{\text{argmax}_j(\tilde{\boldsymbol{Y}}_i) \neq \text{argmax}_j(\hat{\boldsymbol{Y}}_i) \text{ or } \text{argmax}_j(\hat{\boldsymbol{Y}}_i) < 1.01 \times \text{argsort}_j(\hat{\boldsymbol{Y}}_i)(-2)\} \tag{S4}$$

$$L_{\text{s1}} = \sum_{i=1}^{n} L_{\text{s1}}^i = \sum_{i=1}^{n} \frac{1}{2}(L_{\text{rec}}^{\text{self}} + L_{\text{cat}}^{\text{self}} + L_{\text{rec}}^{\text{neigh}} + L_{\text{cat}}^{\text{neigh}}) \tag{S5}$$

$$L_{\text{s2}} = \sum_{i=1}^{n} L_{\text{s2}}^i = \sum_{i=1}^{n} L_{\text{GMVAE}}^i + \text{CE}(\hat{\boldsymbol{Y}}_i, \tilde{\boldsymbol{Y}}_i) \cdot I_{\text{CE}}^i \tag{S6}$$

$$L_{\text{s3}} = \sum_{i=1}^{n} L_{\text{s3}}^i = \sum_{i=1}^{n} L_{\text{GMVAE}}^i \tag{S7}$$

For DLPFC, zebrafish, osmFISH, and STARmap datasets, we used valinna autoencoders with 3, 3, 5, and 3 layers with each layer consisting of 64 neurons to refine the predicted results. For Stereo-seq dataset, we reported the raw results without refinement.

#### S8 Expanding to three dimensions for multi-slice datasets

For multi-slices training of Visium datasets, we constructed the third coordinate as:

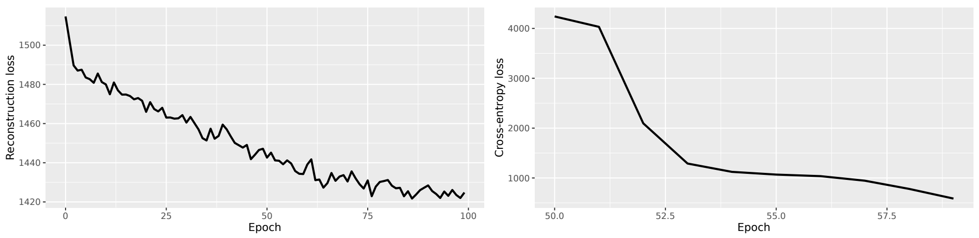


**Supplementary Figure S7**: Reconstruction loss and cross-entropy loss curves on slice 151510 of the DLPFC dataset.

$$s_n \times z_{tech} \times \min_{i,j} (\boldsymbol{C}_i, \boldsymbol{C}_j) \tag{S8}$$

where $s_n$ is the slice index starting from 0, $z_{tech}$ is the expanding factor for the third coordinate and $\boldsymbol{C}_i$ is the spatial coordinates of the $i$-th unit. The rightmost term computes the shortest distance between two units. As the spot organization of Visium is honeycomb-like, $z_{tech}$ is set to 1.5 such that the 6 first-order spatial neighbors of a target unit come from the same slice as the target unit and the 12 second-order spatial neighbors are from two adjacent slices if these slices can be perfectly aligned. For Visium technology, $z_{tech}$ can be set within $\left(1, \sqrt{3}\right)$, where the values of the start point and the end point of this range are the closest pairwise distance and the second closest pairwise distance, respectively.

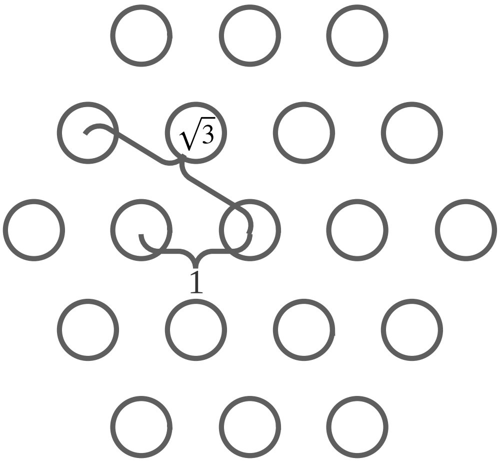


**Supplementary Figure S8**: The expanding factor is set to cover the first-order spatial neighbors but not the second-order spatial neighbors in the same slice.

#### S9 Comparison with other domain clustering methods

*HMRF* HMRF is a probabilistic model designed for identifying spatially variable genes and spatial domains. We used the HMRF method integrated into Giotto [33] and followed the tutorial designed for Visium datasets to predict spatial domains for the Visium DLPFC dataset.

*SpaGCN* SpaGCN is a deep learning model that leverages the power of graph convolutional networks for spatial domain clustering. We followed the tutorial in the official Github repositories to perform spatial domain clustering. Mclust instead of Louvain clustering method was used to ensure the predicted spatial domain numbers were consistent with the ground truth spatial domain numbers.

*BayesSpace* BayesSpace is a probabilistic model that models spatial domains with the Markov random field. We followed their tutorial to perform spatial domain clustering. 2,000 highly variable genes and 20 principal components were used as recommended by the tutorial. BayesSpace might fail on large datasets and we followed the authors' suggestion to divide the whole clustering process of spatialCluster function with the parameter nrep=50,000 into two identical parts with the parameter nrep=25,000 for each.

*SEDR* SEDR is a deep learning model utilizing self-supervised learning for spatial domain clustering. We followed their settings in the tutorial to perform spatial domain clustering. 2,000 highly variable genes and 200 principal components were used as recommended by the tutorial. 12 spatial neighbors were used to construct the adjacency graph for network input. For multi-slice training, we followed their batch integration tutorial to combine slices and graphs constructed from them. The harmony procedure used for batch effect removel is ignored as the batch effect is not obvious in the DLPFC dataset.

*DeepST* DeepST is a deep learning model that combines a graph autoencoder and a denoising autoencoder to extract embedding for spatial domain clustering. We used the default setting in the tutorial for spatial domain clustering. 200 principal components were used to train DeepST for 1,000 epochs.

*BASS* BASS is a Bayesian hierarchical modeling framework for identifying spatial domains. For the STARmap mouse cortex dataset, we used the codes provided in the STARmap tutorial. For datasets generated by other technologies, we followed the tutorial of processing the Visium dataset.

*STAGATE* STAGATE adopts the attention mechanism in convolutional network for spatial domain clustering. For the STARmap mouse cortex dataset, we followed the STARmap tutorial. For datasets generated by other technologies, we followed the tutorial of processing the Visium dataset. Mclust instead of Louvain clustering method was used to ensure the predicted spatial domain numbers were consistent with the ground truth spatial domain numbers. For multi-slice training, we followed their multi-slice without batch effects tutorial.

*GraphST* GraphST employs contrastive learning for better representation to perform spatial domain clustering. We followed the tutorial with default settings to perform spatial domain clustering. Mclust instead of Louvain clustering method was used to ensure the predicted spatial domain numbers were consistent with the ground truth spatial domain numbers. For multi-slice training, we followed their vertical SRT integration tutorial with the PASTE2 aligned anndata file as input.

#### S10 Spatial domain clustering results of the DLPFC dataset on slice 151674 by integrating histology image

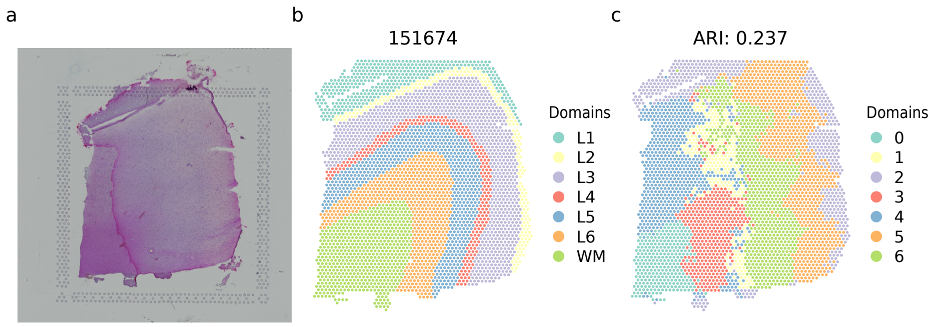


**Supplementary Figure S9**: Visualization of slice 151674. (**A**) Histology image. (**B**) Annotation of spatial domains. (**C**) Predicted spatial domains.
